# Do We See Scale?

**DOI:** 10.1101/371948

**Authors:** Paul Linton

## Abstract

The visual system is supposed to extract distance information from the environment in order to scale the size and distance of objects in the visual scene. The purpose of this article is to challenge this account in three stages: First, I identify three shortcomings of the literature on vergence as our primary cue to near distances. Second, I present the results from two experiments that control for these shortcomings, but at the cost of eradicating vergence and accommodation as effective distance cues (average gain of *y* = 0.161*x* + 38.64). Third, I argue that if all our cues to distance are either (a) ineffective (vergence; accommodation; motion parallax), (b) merely relative (angular size; diplopia), or (c) merely cognitive (familiar size; vertical disparity), then the visual system does not appear to extract absolute distance information, and we should be open to the possibility that vision functions without scale.

## 1 Introduction

Scale has two components: size and distance. Specifically, it is concerned with differentiating a small object up close from a large object far away, even though both may have the same visual angle. Ptolemy (c.160 AD) first articulated visual scale in these terms, arguing that size wasn’t just a function of visual angle (vs. Euclid, c.300 BC), but visual angle appropriately scaled by distance information (see Hatfield, 2002). But the problem is where does the distance information come from? Ptolemy could rely on ‘extramission’: the length of the rays emitted, and then returning, to the eye. As an ‘intromission’ theorist, al-Haytham (c.1021) had no such luxury and relied on familiar size instead. But Descartes (1637) regarded familiar size as a merely cognitive cue and so, along with Kepler (1604), outlined three optical / physiological cues that could plausibly replace Ptolemy’s ‘extramission’ thesis: 1. vergence (the angles of the eyes), 2. accommodation (the curvature of the intraocular lens), and 3. motion parallax (the change in the visual scene from the motion of the observer).

## 2 Vergence

The visual system’s ability to extract distance information from vergence was one of visual psychophysics’ earliest concerns, and was answered affirmatively by Hueck (1838); Meyer (1842); Wheatstone (1852); and Wundt (1862). The contemporary literature is comprehensively reviewed in Appendix A, but suffice to say the effectiveness of vergence as a cue to near distances (up to 2m) is considered settled, and the only real question is whether vergence should be regarded as a veridical distance cue or not? Two papers, Mon-Williams & Tresilian (1999) and Viguier, Clément, & Trotter (2001), are generally cited as authoritative, and their results are summarised in Fig.1.

**Fig.1.** Distance estimates plotted against the vergence demand of the stimulus in Mon-Williams & Tresilian (1999) (left) and Viguier, Clément, & Trotter (2001) (right). © SAGE Publications

Mon-Williams & Tresilian (1999) asked subjects to point with a hidden hand to the distance of a point of light viewed in darkness. The point of light was aligned with the visual axis of the right eye, whilst the vergence demand of the left eye was varied with base-in and base-out prisms. They found a strong linear relationship between the vergence specified distance and the perceived distance of the point of light (*y* = 0.86*x* + 6.5 for distances between 20cm and 60cm). Viguier et al. (2001) presented subjects with a small 0.57° disc in darkness for 5s, then after 5s in complete darkness asked subjects to match a visible reference to the disc’s distance. They found subjects were close to veridical between 20cm and 40cm, but distances were increasingly underestimated beyond that: 60cm was judged to be 50cm, and 80cm judged to be 56cm. Nonetheless, they concluded that their results were consistent with Mon-Williams & Tresilian (1999), at least in reaching space: ‘…the results of our experiment indicate that vergence can be used to reliably evaluate target distance. This is particularly effective in the near visual space corresponding to arm’s length.’

The purpose of this paper is to challenge this conclusion. My concern with these two papers, and indeed with the literature as a whole, is the way the stimuli are presented. Subjects are typically sat in complete darkness with their vergence in a resting state, and then stimuli are suddenly presented at distances as close as 20cm. The sudden presentation of near stimuli necessarily introduces three confounding cues:

1. Diplopia: When the stimuli are initially presented, they are seen as double. But we know that diplopia can be an effective cue to distance. Morrison & Whiteside (1984) found that 90% of performance in estimating the distance of a point of light between 0.5m-9.2m could be attributed to diplopia, since subject performance was only degraded by 10% when the stimuli were shown for 0.1-0.2s, which is too quick for a vergence response.

2. Changing retinal image: An additional cue may be the changing retinal image as the stimulus is converged from its initial diplopic state into a single fused stimulus. Subjects literally watch the two dots (or two discs) approach each other and fuse into one.

3. Kinaesthetic sensation: A final concern is that the subjects are simply be aware of the sudden rotation of their eyes when they are required to immediately converge from their resting state onto a stimulus as close as 20cm. This kinaesthetic cue is directly appreciated, and is quite distinct from vergence as traditionally conceived: it does not need either to (a) feed into, and be processed by, the visual system, nor (b) be presented as visually perceived depth in order to be appreciated. Furthermore, these extreme vergence eye movements have very little ecological validity, and would therefore be a dubious basis upon which to rest the validity of vergence as an egocentric distance cue.

## 3 Experiment 1

Experiment 1 therefore replicates the fundamental aspects of Mon-Williams & Tresilian (1999)’s experimental set-up, whilst controlling for these three confounding cues in two fundamental ways: First, to avoid any sudden jumps in vergence demand, vergence was varied between each trial by having subjects cross-fuse a fixation target, and slowly increasing or decreasing the separation between the target for the left eye and the target for the right eye over 15s. Second, the amount that the vergence demand was varied over these 15s was a modest 1.6º–1.9º, rather than the 17.6º from the sudden presentation of a stimulus 20 cm away.

### Observers

The observers were 12 acquaintances of the author (9 male, 3 female; ages 28-36, average age 31.2) who had indicated their interest in volunteering for the study in response to a Facebook post. All subjects were naïve as to the purpose of the experiment. Observers either did not need visual correction or wore contact lenses (no glasses). All observers gave their written consent, and the study was approved by the School of Health Sciences Research Ethics Committee, City, University of London in accordance with the Declaration of Helsinki. Three preliminary tests were performed to ensure that (a) the subjects’ arm reach was at least 60cm, (b) their convergence response was within normal bounds (18D or above on a Clement Clarke Intl. horizontal prism bar test), and (c) that their stereoacuity was within normal bounds (60 sec of arc or less on a TNO stereo test). 1 additional observer (LN) was excluded as he experienced diplopia from the outset of the experiment.

### Apparatus

Since the purpose of the experiment was to replicate the fundamental aspects of Mon-Williams & Tresilian (1999)’s experimental set-up, a viewing box similar to theirs (45cm high, 28cm wide, and 90cm long) was constructed: see Fig.2. The insides of the box were painted with blackout paint mixed with sand in accordance with the instructions of the optical instrument maker Gerd Neumann Jr (reference in bibliography).

**Fig.2.**
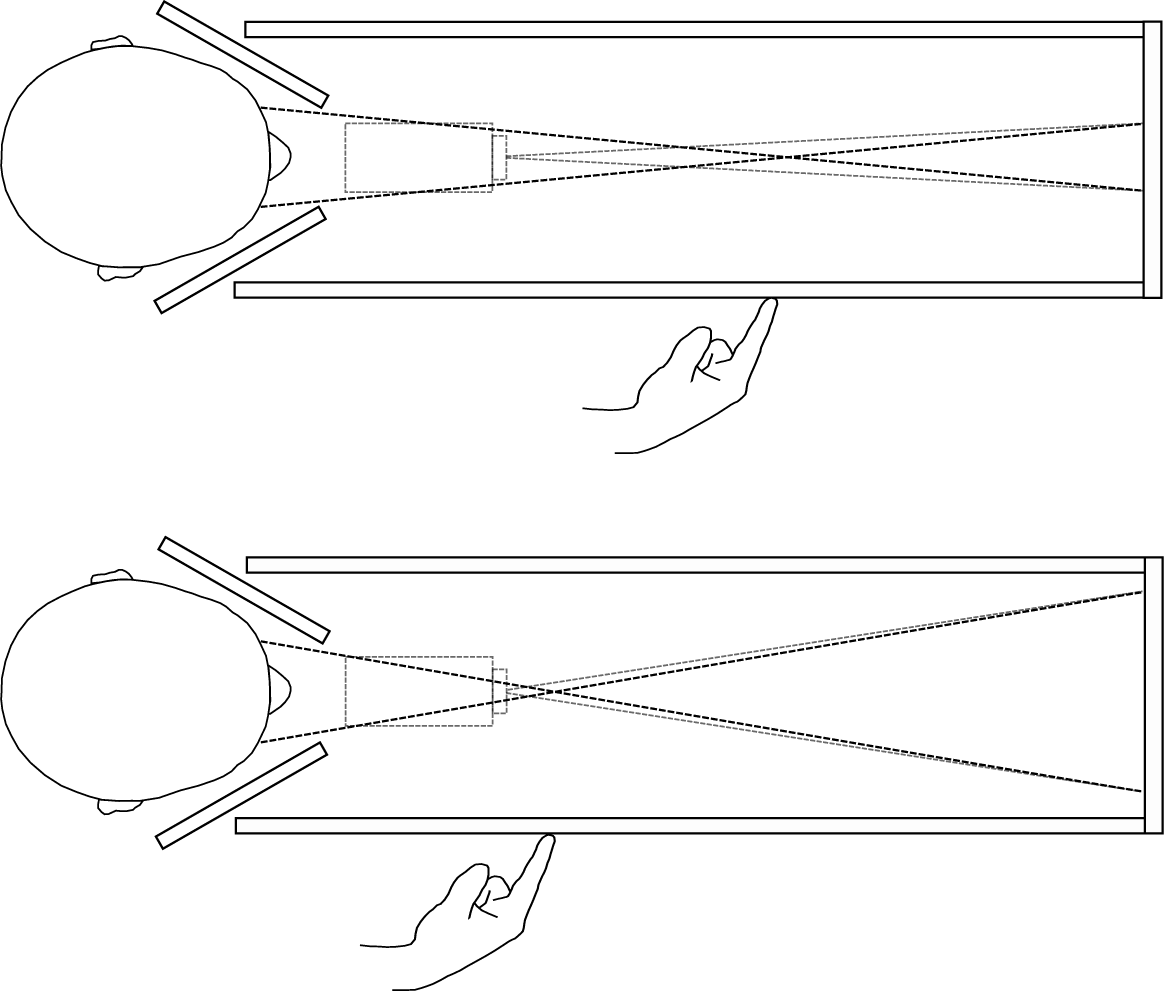
Apparatus for Experiment 1 viewed from above. Laser projector (in grey) projects two stimuli onto a black metal plate at the end of the apparatus. Occluders ensure that the left eye sees only the right stimulus and the right eye sees only the left stimulus (black dotted lines). Vergence angle is varied by increasing the distance between the two stimuli (compare top and bottom images). Subjects point to the perceived distance on a ruler on the side of the box.

I altered Mon-Williams & Tresilian (1999)’s experimental set-up in two fundamental ways:

1. Stimulus alignment: Mon-Williams & Tresilian (1999) aligned their stimulus with the optical axis of the right eye, and varied the vergence demand of the left eye using prisms. This approach has two shortcomings: First, it leads to an asymmetric vergence demand. For a 20cm target aligned with the right eye, the vergence demand is 17.75° for the left eye and 0° for the right eye, rather than 8.81° for each eye had the target been located along the midline. In normal viewing conditions head rotations would eradicate such extreme asymmetries in vergence demand. Second, the stimulus is liable to be perceived as drifting rightwards as it gets closer: at 50cm a stimulus aligned with the right eye is offset from the subject’s midline by 3.5°, whilst at 20cm it is offset by 8.8°. I therefore aligned the stimulus with the subject’s midline.

2. Stimulus presentation: As I have already discussed, rather than varying the vergence demand with prisms, which introduce sudden jumps in vergence, I had subjects cross-fuse a fixation target and varied vergence by slowly increasing or decreasing the separation between the targets for the left and right eyes. Occlusion barriers were used to ensure each eye only saw its appropriate stimulus. The stimuli were presented using a Sony MP-CL1A laser projector, fixed 25cm in front and 5cm below the line of sight, which projected the stimuli onto a black metal plate that formed the back wall of the apparatus. Laser projection was used to ensure the stimuli were viewed in perfect darkness (unlike CRT or OLED displays, lasers projectors emit no light for black pixels, ensuring perfect black values and no residual luminance).

Subjects were completely naïve about the experimental set-up: (a) they did not see the room or apparatus beforehand, which was sealed off by a curtain, and (b) they were wheeled into the experimental room wearing a blindfold, their hand was guided to a head and chin rest, and they had to ensure their head was in place, with a further hood of blackout fabric pulled over their head, before they could take the blindfold off. This procedure, coupled with the fact that the box was sealed, and the illumination in the room outside the box was reduced to a low-powered LED, ensured subjects viewed an isolated stimulus in perfect darkness.

Before the start of each trial subjects were asked to indicate how many targets they could see, and the metal occluders were adjusted by the experimenter until only one target was visible in each eye. The experimenter then guided their arm to a ruler, attached at eye level on the right-hand side of the box (offset to the right of the subject’s midline by 12cm), and subjects were told to indicate their distance judgements on the ruler, but to relax their arm by their sides between trials. Distance judgements were recorded by the experimenter.

The stimuli, also presented at eye level, were produced in PsychoPy (Peirce, 2007; 2009). They comprised of (a) a fixation target (300 green dots located within a 1.83° circle, with the dots randomly relocating every 50ms within the circle, giving the impression of a disc of shimmering dots): see Fig.3, and (b) a single green dot that subjects had to point to. The fixation target changed in size (sinusoidally between 1.83° and 0.91° at 1Hz). The shimmering and constantly changing size of the fixation target ensured that as the vergence demand was varied, any residual motion-in-depth from retinal slip would be hard to detect.

**Fig.3.**
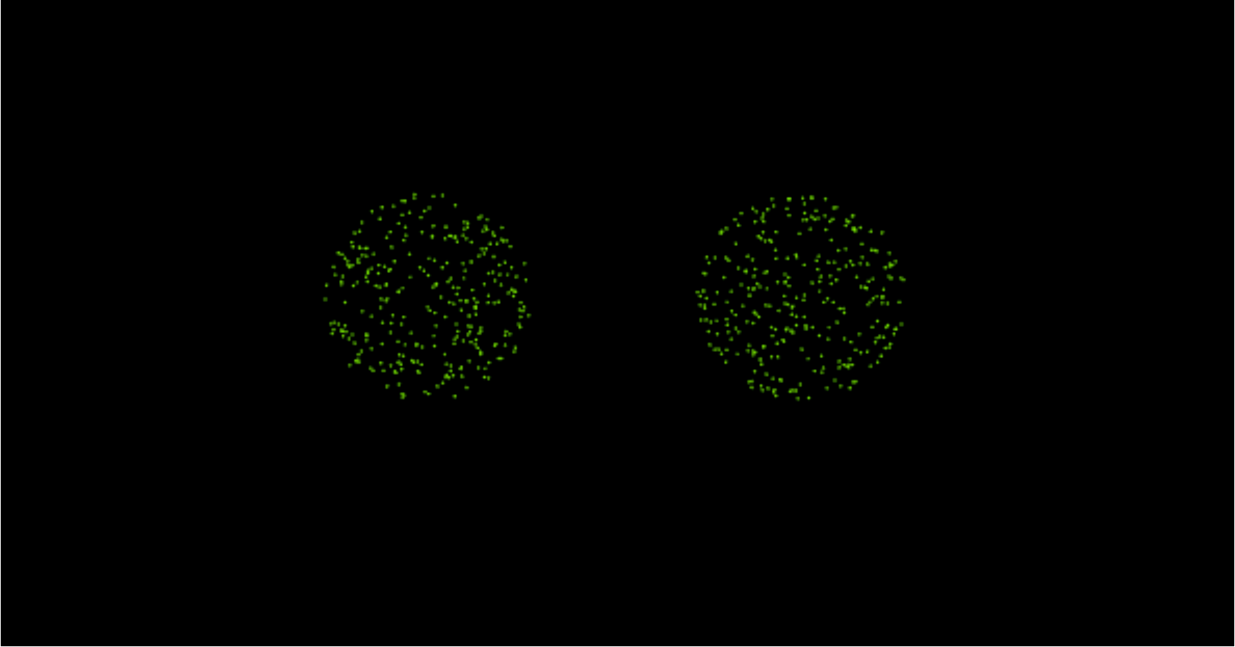
The fixation target in Experiment 1, comprising of 300 dots with their location within the circle updating randomly every 50ms.

Subjects completed 4 sets of 24 trials, with 96 trials in total per observer. The initial stimulus was the fixation target with the vergence angle specified for 50cm. Once subjects confirmed that the stimulus was fused, the target changed in size sinusoidally between 1.83° and 0.91° at 1Hz for 32s. Unbeknownst to the subjects, their vergence angle was also slowly increased from 50cm to 29cm. The stimulus then changed to a dot and subjects had to indicate its distance by pointing on the side of the box. At this point the stimulus changed back to the fixation target for 15s, and vergence was stepped up or down for the next trial using a pseudo-random walk. The pseudo-random walk covered 7 distances: 20cm, 22cm, 25cm, 29cm, 34cm, 40cm, 50cm and specified the vergence distance for the remaining 23 trials of each set. The vergence demand of the next trial was either stepped up or stepped down by one step. The walk was only pseudo-random as there was a slightly higher probability (0.6) that the step would be away from the middle of the range, to ensure full coverage of the 7 distances. Walks were simulated prior to the experiment, and a walk was chosen for each subject that ensured each of the 7 distances was tested at least 10 times over the 96 trials.

After the main experiment was completed, a control study was run in full-cue conditions to confirm Mon-Williams & Tresilian (1999)’s and Swan et al. (2015)’s findings that manual pointing with a hidden hand is a good proxy for perceived distance. The control replicated the head and chin rest, right-hand wall, and hidden ruler of the original apparatus in Experiment 1, but removed the top, back, and left-hand side of the box, enabling a familiar object (a 510g Kellogg’s Rice Krispies box) to be seen in full-cue conditions. Subjects pointed to the front of the cereal box with a hidden hand in 3 sets of trials that ensured 10 trials in total for each of the 7 distances (20cm, 22cm, 25cm, 29cm, 34cm, 40cm, and 50cm). One subject (SM) was unable to return to complete the control.

### Results

The results of the main experiment and the control are shown in Fig.4, with the results of the main experiment in black and the results of the control in grey.

**Fig.4.**
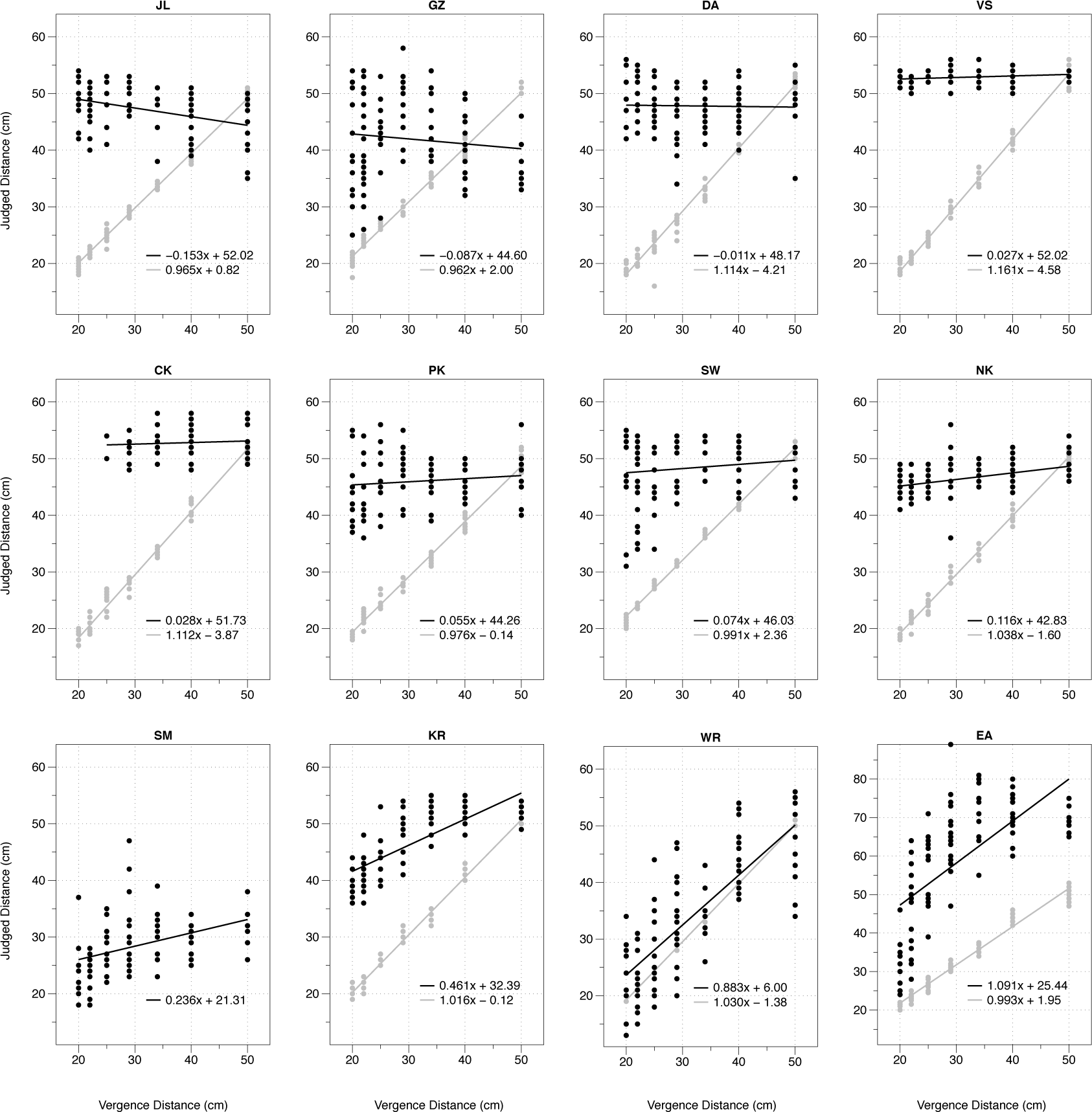
Results of Experiment 1. Grey dots show the results for the full-cue pointing control. Black dots show the results for the vergence-only condition.

In the control, all subjects were close to veridical when it came to pointing with a hidden hand at the distance of a familiar object in full-cue conditions. A linear mixed effects model (Pinheiro & Bates, 2000) conducted in R (R Core Team, 2012) using the lme4 package (Bates, Maechler, & Bolker, 2012; see Lisi, 2015) estimates the relationship as *y* = 1.032*x –* 0.76 (with 95% confidence intervals of 0.992 to 1.071 for the slope, and –2.36 to 0.73 for the intercept). This result confirms the findings of the control tests in Mon-Williams & Tresilian (1999) (*y* = 1.08*x –* 1.35) and Swan, Singh, & Ellis (2015) (*y* = 1.005*x –* 2.76) that hidden hand pointing is an accurate way of reporting perceived distance.

Turning to the results of the main experiment, we find significant individual differences in the effectiveness of vergence as a distance cue: 8 subjects with no gain whatsoever, 2 subjects with very modest gains (SM: 0.24; KR: 0.46), and 2 subjects with very significant gains (WR: 0.88; EA: 1.1, although EA consistently overshoots by 25cm). To try and make sense of these individual differences, we can cluster the histogram of the slopes in Fig.4 using a Gaussian mixture model in R with the mclust5 package (Scrucca, Fop, Murphy, & Raftery, 2017). According to the Bayesian information criterion (BIC), a model with two populations and equal variance best fits the data: one population with 10 subjects and an average slope of 0.074, and another population with 2 subjects and an average slope of 0.983: see Fig.5. We can also analyse these two populations using a linear mixed effects model: the first population of 10 subjects has a relationship of *y* = 0.075*x* + 43.52 (with 95% confidence intervals of –0.035 to 0.183 for the slope, and 37.12 to 49.70 for the intercept), and the second population of 2 subjects has a relationship of *y* = 0.987*x* + 15.72 (with 95% confidence intervals of 0.747 to 1.219 for the slope, and –3.94 to 36.13 for the intercept).

**Fig.5.**
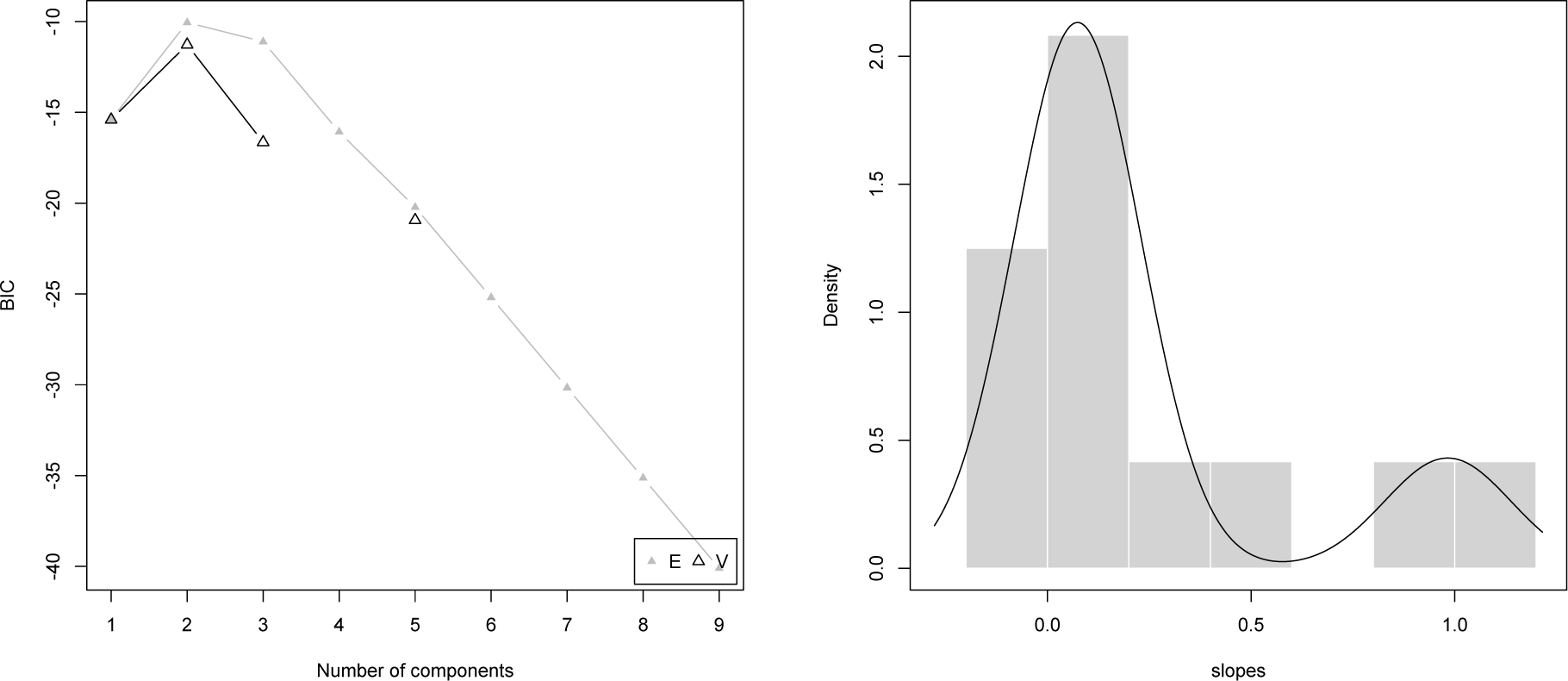
The Bayesian Information Criterion (BIC) indicates that two groups of equal variance best fit the histogram of the slopes in Fig.4 (left), with the resulting Gaussian distributions plotted on top of the histogram of the slopes from Fig.4 (right).

### Discussion

The general finding of Experiment 1 is that vergence is an ineffective distance cue for the vast majority of the population, with an average gain of *y* = 0.075*x* + 43.52 for 10 of the subjects. This suggests that the high gains reported in Mon-Williams & Tresilian (1999) (*y* = 0.86*x* + 6.5 between 20cm and 60cm) and Viguier et al. (2001) (close to veridical between 20cm and 40cm) can be attributed to the three confounding cues I identified in Section 2: (1) diplopia, (2) the changing retinal image, and (3) vergence as a kinesthetic cue.

But we still have to explain why the two outliers (WR and EA) have a slope close to 1? On the face of it, the stark contrast between 10 subjects with a slope close to 0, and 2 subjects with a slope close to 1, might appear to suggest that vergence is an effective distance cue for a small minority of the population. But I believe this conclusion is premature, since all three of the subjects with the highest gains (KR, WR, and EA) made comments during the experiment and/or during their post-experiment debrief that were consistent with vergence / accommodation conflict accounting for their results. The problem is that Experiment 1 induced up to 3.9D of vergence / accommodation conflict, with subjects accommodating on a back plane 90cm away, but converging to distances as close as 20cm. This conflict can manifest itself in three different ways:

First, accommodation could determine vergence, and break binocular fusion. This was only a concern for one subject (CK), who could fuse the fixation target at the closest distances (20cm, 22cm), but whose fusion broke as soon as the dot was presented. Two other subjects experience diplopia (JL twice, GZ once).

Second, vergence could determine accommodation, and the dot could go out of focus. Two of the subjects with the highest gains (KR and EA) reported this, and both reported using the change in size as the dot went out of focus as a distance cue.

Third, accurate vergence and accommodation could be maintained but at the cost, for some subjects, of significant eye strain. In a preliminary test, two very experienced psychophysical observers reported eye strain, and WR, the subject with the second highest gain, also reported eyestrain, describing the experiment as ‘exhausting’ for his eyes.

This suggests that the high gains reported by KR, WR, and EA may be explained by vergence / accommodation conflict rather than the effectiveness of vergence as a distance cue. The solution is to retest these three subjects with correct accommodation cues in place. But this poses a challenge: if vergence is being varied dynamically, how can we vary accommodation dynamically so it is kept in step with vergence?

One solution is to avoid this problem altogether by keeping both vergence and accommodation fixed and consistent at the same target distance (apart from the initial transition of the fixation target from its 50cm starting point to the first target distance), and to test 10 consecutive trials at that distance, with the accommodation cues appropriately controlled for using trial lenses. This method was attempted with WR, but soon abandoned: He was tested with vergence and accommodation specifying 20cm. Whilst he could maintain fixation on the fixation target, as soon as it turned into a dot he found it very difficult to maintain focus and asked to stop the experiment after 6 trials. In his own words, the eye strain was ‘painful’, ‘quite exhausting for his eyes’, and ‘much worse’ than before. Although there was no vergence / accommodation conflict at the point at which the dot appeared, the experimental set-up appears to have induced the illusion of such a conflict. One explanation is that whilst the slightly faint and indistinct fixation target of shimmering dots may facilitate open-loop accommodation, a single static dot does not. Consequently, when the dot was presented, a sudden sharp accommodative response might have been required, without any attendant change in vergence, leading to the illusion of vergence / accommodation conflict.

WR was no longer available to continue with the study. A new paradigm was adopted for KR and EA. But before this second experiment is introduced it is worth considering accommodation as a distance cue in its own right.

## 4 Accommodation

Accommodation is regarded as a less reliable distance cue than vergence. Fisher & Ciuffreda (1988) had subjects point to the distance of a target viewed in darkness in a Badal system (to control for size and luminence), and found a relationship of *y* = 0.27*x* + 2.33 for 16cm-50cm. However, this low average gain masked a high degree of individual and inter-stimulus differences: individual gains ranged from 0 to 0.8 for a cross stimulus, and –0.20 to 0.92 for a Snellen chart. Mon-Williams & Tresilian (2000) found similar results (varying stimulus size between trials, rather than using a Badal system), but unlike Fisher & Ciuffreda (1988) concluded that accommodation is not an absolute distance cue because even when they focused on their two observers with the highest gains (*y* = 0.62*x* + 1.55 and *y* = 0.60*x* + 0.88 for 16cm-40cm), they argued that the variability in their responses was so great that ‘it is clear that accommodation is providing no functionally useful metric distance information for these observers. The responses were unrelated to the actual distance of the target.’

Mon-Williams & Tresilian (2000) instead suggest that subjects may only be able to extract ordinal depth information from accommodation (i.e. whether the present trial is closer or further away than the previous one). But this doesn’t explain how two subjects could have the same ordinal depth success rate (78-79%) but completely different gains (*y* = 0.62*x* + 1.55 vs. *y* = 0.28*x* + 1.49), nor the fact that when Liu, Hua, & Cheng (2010) presented subjects with a monocularly viewed half Siemens star at three possible accommodative distances (20cm, 33cm, 50cm) the average success rate was 80% (with individual performance ranging from 67% to 93%), even though if the previous distance were 20cm or 50cm, ordinal depth would provide no information by which to choose between the other two alternatives.

Subjects therefore appear to be able to extract some useful metric distance information from accommodation. But I have two concerns:

First, as I explain in Linton (2017), p.119, rather than responding to accommodation as a physiological cue to distance, the visual system may simply be responding to any number of optical changes in the retinal image that correlate with increased accommodation. For instance, increases in (a) monochromatic aberrations, (b) chromatic aberrations, or (c) low-frequency microfluctuations. By contrast, Pentland (1987) assumes a fixed chromatic aberration (of 1D), and a fixed microfluctuation rate (of 2Hz, from high-frequency microfluctuations), to suggest that the visual system may simply be responding to the difference in defocus blur between two retinal images, either (a) two sequential images (from microfluctuations), or (b) the images from two different photoreceptors (from chromatic aberration).

Second, the stimulus presentation in Fisher & Ciuffreda (1988), Mon-Williams & Tresilian (2000), and Liu et al. (2010), mirrors my concern with the stimulus presentation in Mon-Williams & Tresilian (1999) and Viguier et al. (2001). Whilst Mon-Williams & Tresilian (1999) and Viguier et al. (2001) study diplopia-driven vergence, Fisher & Ciuffreda (1988), Mon-Williams & Tresilian (2000), and Liu et al. (2010) study blur-driven accommodation. This brings with it 3 equivalent confounding cues: (1) the initial blur in the stimulus, (2) the changing retinal image as the stimulus is brought into focus, and (3) large sudden changes in accommodation as a kinaesthetic sensation.

One way to control for these three confounding cues is to vary the accommodative demand gradually rather than suddenly, much like my gradual variation of vergence in Experiment 1. This can be achieved by slowly moving a stimulus back and forth behind a Badal lens which controls for angular size and luminance. As I report in Linton (2017), p.132, moving a target back and forth behind a Badal lens appears to have no effect on its perceived distance. Indeed, unlike the equivalent observation for vergence (Regan, Erkelens, & Collewijn, 1986), this appears to hold true for a single dot as well. It therefore appears that accommodation does not provide an effective cue to distance once defocus blur is eliminated.

But Badal setups prove notoriously difficult for most subjects (see Charman & Heron, 2015), with only 16% of the subjects in Metlapally, Tong, Tahir, & Schor (2014), and 20% of the subjects in Metlapally, Tong, Tahir, & Schor (2016), able to accommodate effectively. So, rather than trying to isolate accommodation as an individual cue, a more effective approach is to incorporate accommodation alongside vergence, and test their effectiveness in combination.

## 5 Experiment 2

Experiment 2 uses trial lenses to incorporate approximately consistent accommodation cues into the experimental set-up of Experiment 1, whilst exploiting the ‘zone of clear single binocular vision’ (the ±1D of accommodation / vergence conflict that observers can reasonably tolerate: see Hoffman et al., 2008; Fig.6). This enables 5 vergence distances (23cm, 26cm, 30cm, 36.5cm, 45.5cm) to be tested using 3 sets of trial lenses that maintain vergence / accommodation conflict within reasonable bounds (as summarised in Fig.6):

1. Near (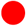) = –4.15D (24cm) to test 23cm, 26cm, and 30cm vergence.
2. Middle (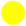) = –3.15D (32cm) to test 23cm, 26cm, 30cm, 36.5cm, and 45.5cm vergence.
3. Far (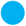) = –2.15D (47cm) to test 30cm, 36.5cm, and 45.5cm vergence.

**Fig.6.**
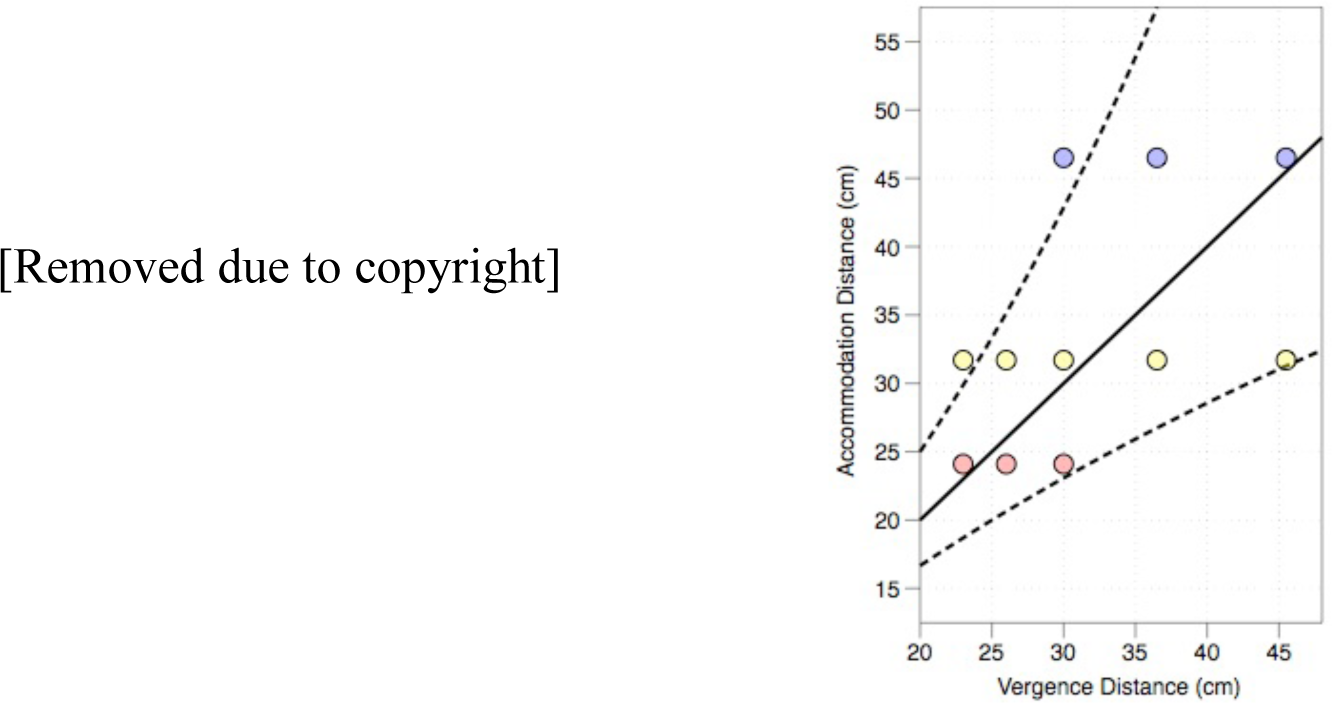
Left: [Removed due to copyright] Graph from Hoffman et al. (2008) illustrating the ‘zone of clear single binocular vision’ (±1D). © ARVO. Right: The various combinations of vergence and accommodation tested in Experiment 2 using trial lenses. 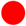 = –4.15D, 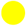 = – 3.15D, and 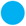 = –2.15D. The solid black line indicates no vergence / accommodation conflict, and the two dotted lines mark the ‘zone of clear single binocular vision’ (±1D of vergence / accommodation conflict).

To improve the accommodative response in light of WR’s apparent difficulties, the fixation target was amended to a high-contrast random-dot 2.4° x 2.4° square, with a bulls’ eye added to the centre of the target after the first two subjects: see Fig.7 (only NM and KL used the original target; its addition is explained in the Discussion).

**Fig.7.**
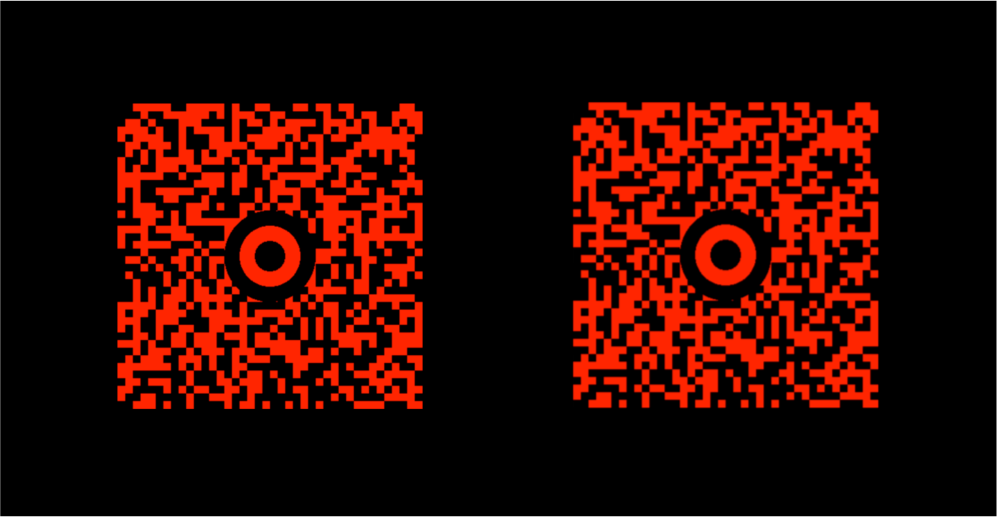
The fixation target in Experiment 1, comprising of two random-dot stereograms with a bulls-eye at the centre of each to facilitate fixation.

As with Experiment 1, the experiment started with subjects fixating on the fixation target. Rather than changing in angular size, the fixation target varied in luminance (between 100% and 50% of its initial luminance, at 2Hz) for 50s. During this period vergence was stepped up from its initial 50cm to the middle of the specific lens’ range (26cm for Near, 30cm for Middle, and 36.5cm for Far). After the 50s, the fixation target turned into a dot and subjects had to point to its distance. As with Experiment 1, the subsequent trials continued in a pseudo-random walk, although the fixation target was visible for 30s between trials rather than 15s.

To increase contrast, the stimuli were projected onto a white screen 156cm away from the observer, rather than a black metal plate 90cm away. To ensure the increase in illumination did not also illuminate the apparatus, black fabric was added to ensure a narrow viewing window, and red filters from red-cyan stereo-glasses (blocking ≈100% green light, ≈90% blue light) were added in front of each eye. In a separate preliminary experiment using an autorefractor, the filters were found to have no impact on accommodation.

The observers were two of the observers from Experiment 1 with high gains (KR and EA) and 12 City, University of London undergraduate students (8 female, 4 male; age range 18-27, average age 20.8) recruited through flyers and Facebook Posts. All subjects were naïve as to the purpose of the experiment. The same exclusion criteria as Experiment 1 were applied, with the additional requirement that the subject’s accommodative response was within normal bounds (tested using an RAF near-point rule). The study was approved by the School of Health Sciences Research Ethics Committee, City, University of London in accordance with the Declaration of Helsinki, and all subjects gave their written consent. The undergraduate students were paid £10/hr + a £20 completion bonus. 6 additional subjects had to be excluded: 4 subjects (MG, HV, FP, NS) because their prism bar convergence was below 18D, and 2 subjects (BB, RE2) because they experienced diplopia from the outset of the experiment.

The 12 undergraduates each participated in 7 sets of 20 trials (2 Near, 3 Middle, 2 Far) in random order, ensuring each lens / vergence combination was tested at least 10 times. KR and EA participated in a reduced version of this experiment with 4 sets of 20 trials (1 Near, 2 Middle, 1 Far), with only 23cm and 26cm tested in the Near condition, 26cm, 30cm, 36.5cm tested in the Middle condition, and 36.5cm and 45.5cm tested in the Far condition.

### Results

The results of the two prior subjects (KR and EA) are plotted in Fig.8 against their previous performance in Experiment 1.

**Fig.8.**
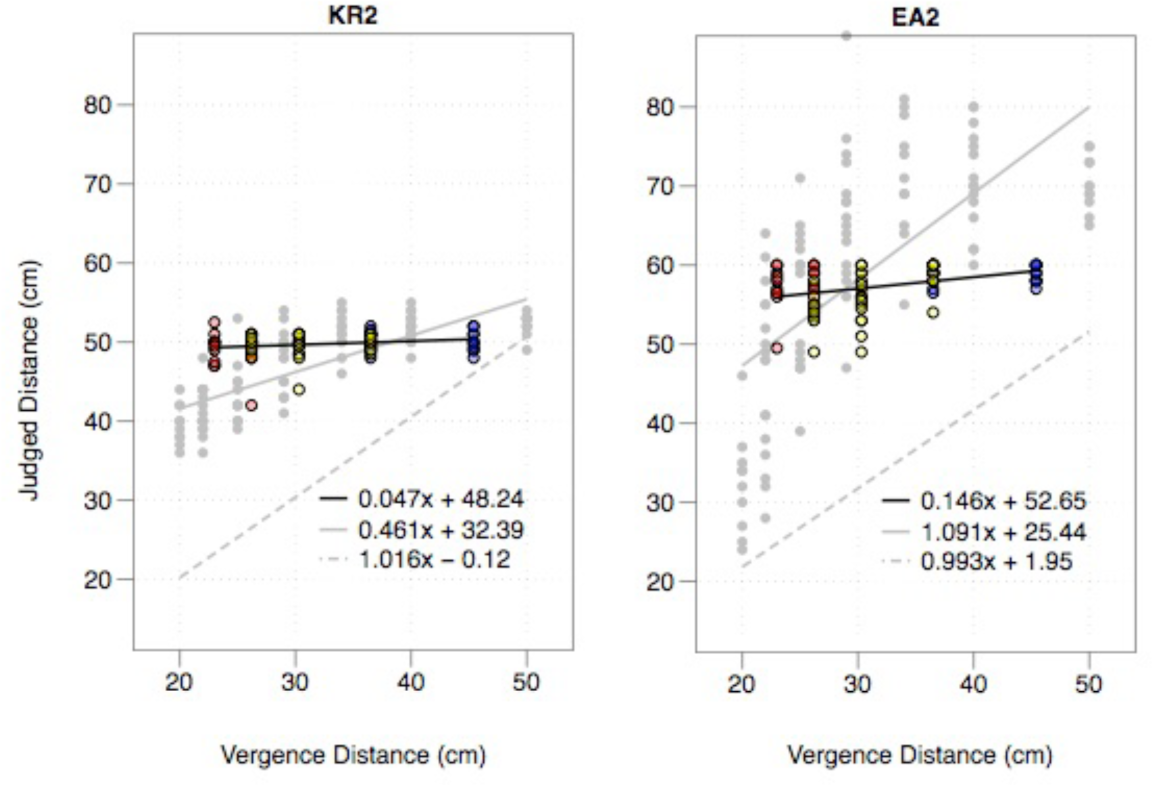
Performance of subjects KR and EA from Experiment 1 compared with their performance in Experiment 2. Performance in Experiment 1 indicated by grey dots and grey line. Performance in Experiment 2 indicated by coloured dots (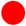 = –4.15D, 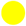 = –3.15D, and 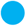 = –2.15D) and black line. Performance in full-cue control condition indicated by broken grey line.

What is striking is how their previously high gains are eradicated with the addition of accommodation cues: KR’s previous performance of *y* = 0.461*x* + 32.39 drops to *y* = 0.047*x* + 48.24, whilst EA’s previous performance of *y* = 1.091*x* + 25.44 drops to *y* = 0.146*x* + 52.65 (both of these drops in performance are significant to p < 0.001). This confirms the hypothesis that the high gains in Experiment 1 were driven by vergence / accommodation conflict, since eliminating this conflict effectively eliminates their capacity to estimate distance. It also confirms the almost complete ineffectiveness of vergence and accommodation as distance cues for these two observers.

The results for the 12 undergraduates are plotted in Fig.9.

**Fig.9.**
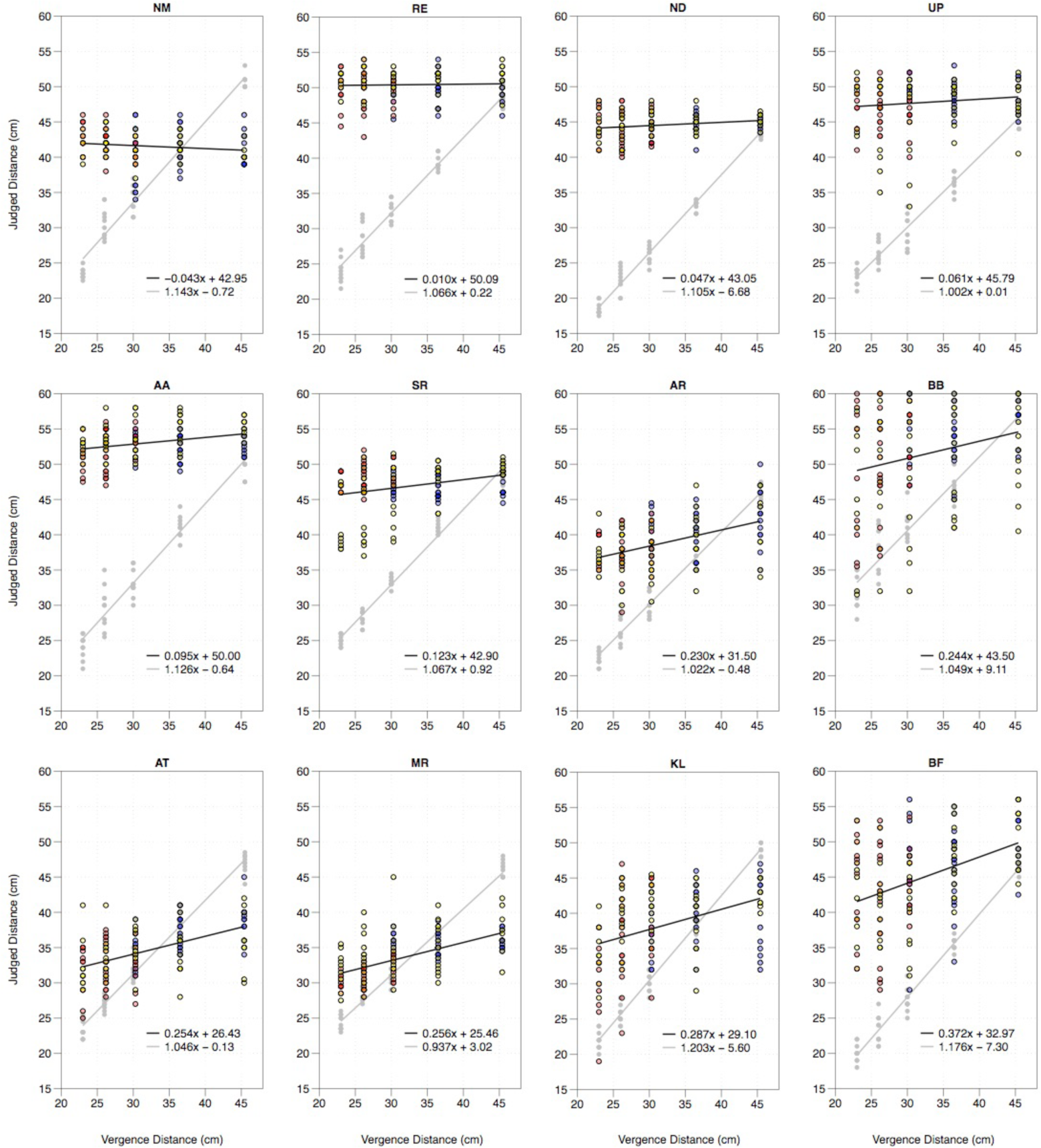
Performance of the 12 new subjects in Experiment 2 indicated by coloured dots and black line. Colour of dots correspond to accommodative demand (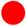 = −4.15D, 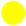 = −3.15D, and 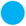 = −2.15D). Grey dots and grey line indicate performance in the full-cue control condition.

Turning to the full-cue control task, we again find a strong linear relationship *y* = 1.078*x –* 0.69 using a linear mixed effects model across the 12 observers (95% confidence intervals of 1.036 to 1.122 for the slope, and –3.19 to 1.81 for the intercept), further confirming hidden hand pointing as an effective reporting mechanism for perceived distance.

But turning to the main experiment, we again find significant individual differences: 6 subjects with virtually no gain, and 6 subjects with a gain of between 0.23 and 0.37. To try and make sense of these individual differences, we can cluster the histogram of the slopes in Fig.9 using a Gaussian mixture model. According to the Bayesian information criterion (BIC) we find a single population with an average gain of 0.161 best fits the data: see Fig.10, although this is only marginally better than two populations with equal variance. Fitting the data to a single population with a linear mixed effects model we find a relationship of *y* = 0.161*x* + 38.64 (with 95% confidence intervals of 0.090 to 0.239 for the slope, and 33.43 to 43.36 for the intercept). To put this value in context, a combined gain of 0.16 from vergence and accommodation is less than 60% of the 0.27 gain that is commonly attributed to accommodation alone as a distance cue (Fisher & Ciuffreda, 1988). Since accommodation is considered an ineffective distance cue, the same must now be concluded of vergence and accommodation in combination.

**Fig.10.**
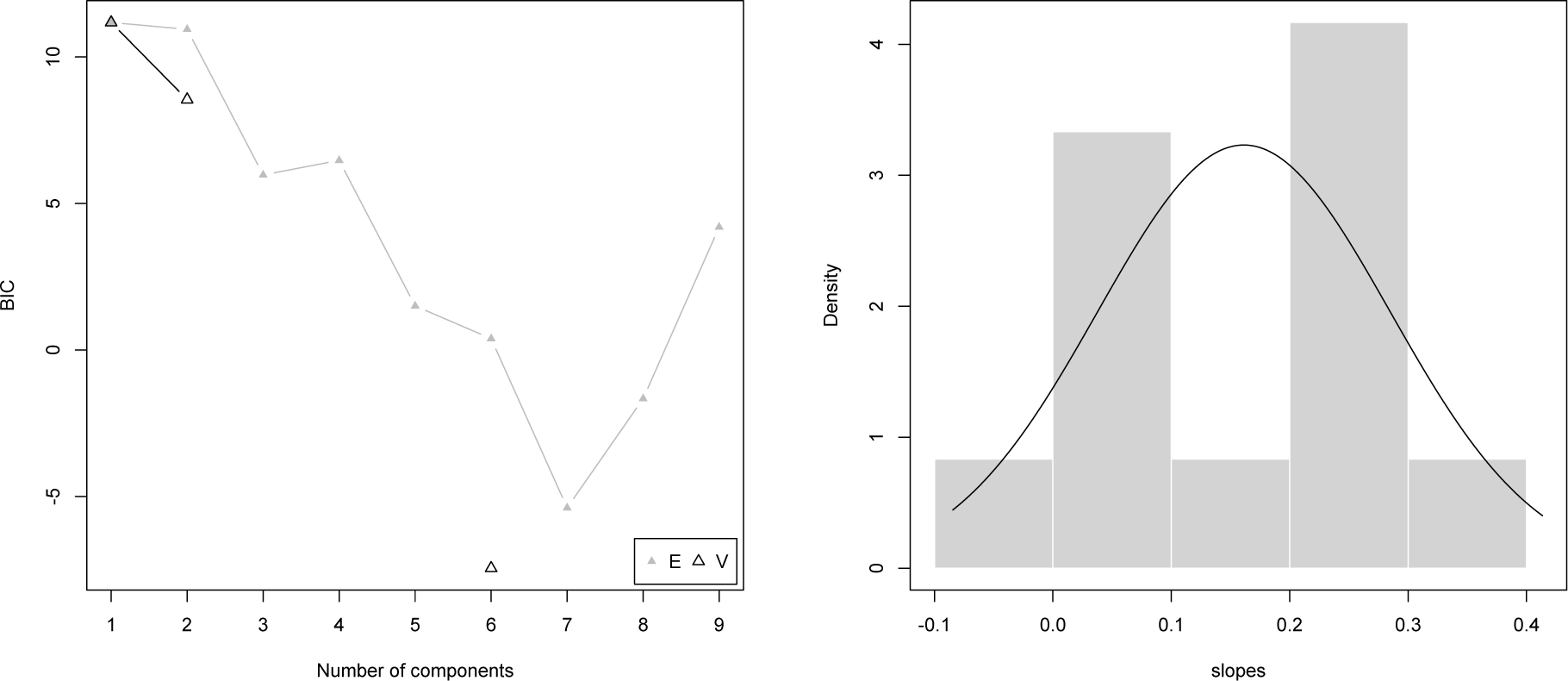
The Bayesian Information Criterion (BIC) indicates that a single population best fits the histogram of the slopes in Fig.9 (left), with the resulting Gaussian distribution plotted on top of the histogram of the slopes from Fig.9 (right).

This conclusion is further supported by the variance of the 6 subjects with the highest gains. We can estimate the standard deviation of the residual error (i.e. how much each subject departs from their own line of best fit in Fig.9), after correcting for motor error (assuming that perceptual error and motor error are independent) using the following formula:

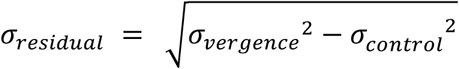

Even if we limit ourselves to ± 2 standard deviations from the slope of best fit to rule out any outliers, we still find residual errors of 27cm for BB, 23cm for BF, 20cm for KL, 12cm for AR, 11cm for AT, and 9cm for MR, with an average of 17cm. Given the range of the experiment itself was only 22.5cm, and pointing was confined to reaching space, one is left questioning just how functionally useful cues with this degree variance could be. Mon-Williams & Tresilian (2000) come to the same conclusion in the context of accommodation alone; although the variance was higher in that study, so too were the average gains.

Finally, Experiment 2 could have been conducted with just the ‘middle’ lens (–3.15D), since this lens covers the full range of vergence distances (see Fig.6). Did varying accommodation by 2 dioptres using the ‘near’ and ‘far’ lens have any effect? And would the results have been any different if we eradicated the ±1D of vergence / accommodation conflict completely? We can test this hypothesis by comparing the results for the 23cm, 30cm, and 45.5cm vergence distances using (a) the ‘middle’ lens for all three of these vergence distances, against (b) the ‘near’ lens for 23cm, the ‘middle lens’ for 30cm, and the ‘far’ lens for 45.5cm, where there is no vergence / accommodation conflict. We only find a marginal reduction in performance from (a) *y* = 0.176*x* + 38.02 when only the ‘middle’ lens is used, to (b) *y* = 0.147*x* + 38.91 when the vergence / accommodation conflict is eradicated, and this reduction in performance is not statistically significant.

This result has two implications: First, the performance in Experiment 2 would have been no better if the vergence / accommodation conflict had been completely eradicated, indeed it might have been marginally worse. Second, the modest gain of 0.16 cannot simply be attributed to the residual vergence / accommodation conflict in the apparatus.

### Discussion

Vergence and accomodation appear to be ineffective cues to distance, even when tested in combination. This is undoubtedly true for half the subjects, for whom there virtually no gain (*y* = 0.049*x* + 45.80 in a linear mixed effects model). However, it is worth considering how the other half were able to achieve modest gains of 0.23–0.37 (*y* = 0.274*x* + 31.49 in a linear mixed effects model)? A number of possible explanations were suggested by the subjects themselves, either during the experiment or in their post-experimental debrief:

1. Eyestrain as kinesthetic and/or proprioceptive cue: Consider BF, the subject with the highest gain. She had 30% more gain than the next highest subject (0.372 vs. 0.287). During the experiment she mentioned that it was ‘messing up my accommodation’, so I asked her to explain this afterwards: ‘It was very clear, but I could feel my eye are working, my eyes are focusing then relaxing then focusing.’ Discussing some of the trials she reported: ‘I really had to focus to stop them going two … the target started to separate when I didn’t really focus on it.’ She described it in the following terms: ‘I usually get the same sensation when I’m up too late and doing some studies – a slight strain in the eye, it’s not too bad, it’s just that you really have to focus.’ Similarly, KL, the subject with the second highest gain (0.287), reported that with near targets she felt her ‘eyes accommodating a lot to get them to work.’

In her description, BF runs two concerns together: one *kinesthetic*, the other *proprioceptive*: The *kinesthetic* cue is the feeling of our eyes working as they struggle to maintain focus and fusion. The *proprioceptive* cue is the discomfort that is often the consequence of these efforts. Note that both are quite distinct from *visual* cues. For instance, the *proprioceptive ‘*Percival’s zone of comfort’ (±0.5D of vergence / accommodation conflict) is narrower than the *visual ‘*zone of clear single binocular vision’ (±1D of vergence / accommodation conflict): see Hoffman et al. (2008) (Fig.6). Since vergence / accommodation conflict appears to affect younger subjects more (see Yang et al., 2012; cf. Read & Bohr, 2014), this might appear to explain why more subjects in Experiment 2 (average age 20.8) reported non-negligible gains than in Experiment 1 (average age 31.2).

However, as we have already noted, we should be cautious of suggesting that if the residual ±1D of vergence / accommodation conflict were eradicated, these gains would disappear. Although the experimental paradigm could be improved by either (a) eradicating the vergence / accommodation conflict altogether (using Optotune electrically tunable lenses), or (b) implementing open-loop accommodation (using Koryoeyetech pinhole contact lenses), I am sceptical that eyestrain can be eradicated from the apparatus completely, even though subjects do not experience it when fixating on a near object in the real world:

First, this is not a natural task: Even though Experiment 2 kept within the ‘zone of clear single binocular vision’, two subjects (BB and RE) had to be excluded because they experienced diplopia from the outset, whilst one subject (SR) experienced diplopia midway through the experiment, and her remaining trials had to be rescheduled. As we have already noted, subjects struggle to accommodate accurately in a Badal system when accommodation is manipulated merely as an optical reflex (Charman & Heron, 2015), and there is increasing evidence that cognitive cues to distance are required for effective accommodation (Gwiazda, Thorn, Bauer, & Held, 1993; Otero, Aldaba, Martínez-Navarro, & Pujol, 2017). The same may well be true when vergence and accommodation are manipulated using purely optical cues.

Second, pointing to a dot in perfect darkness is the visual equivalent of an echo chamber. You are aware of physical sensations that you wouldn’t necessarily notice in natural viewing conditions. If subjects had nothing else to go on, they are liable to give weak physical sensations much more weight than they ordinarily would.

2. Retinal slip: KL, the subject with second highest gain (0.287), reported that often the dot seemed to ‘jump in front or behind the fixation plane’ when it appeared: ‘sometimes it appeared to shoot out the front, sometimes it appeared to shoot out the back.’ This would appear to suggest that her vergence slightly lagged behind the changing vergence of the fixation target, and then suddenly corrected when the dot was presented. A bulls’ eye was subsequently added to the fixation target to facilitate vergence. BB, another subject with a higher gain (0.244), also described the dot as ‘jumping forward, sometimes jumping back … from the fixation plane.’

3. Vergence-hand-proprioception linkage: MR, the subject with the third highest gain (0.256) suggested in her debrief that the dot always looked the same distance away. Instead, she would put her hand out and converge back and forth between the visible dot and her hidden hand until she felt they were at the same distance. Whilst no-one else explicitly adopted this strategy, AT, the subject with the fourth highest gain (0.254), suggested that she was unable to judge the distance of the dot until she had put her hand out, whilst BF made a similar comment after one of the trials: ‘It looks far away and I put my hand out, but when I put my hand out I realise that it’s closer.’ For discussions of our ability to converge onto a hidden hand see: Erkelens, Steen, Steinman, & Collewijn (1989); Koken & Erkelens (1992); Koken & Erkelens (1993); Mon-Williams, Tresilian, Plooy, Wann, & Broerse (1997).

4. Vergence-action linkage: Swenson (1932) reported that ‘even in those occasional judgments where the subject reported a ‘lost’ feeling, where he felt his judgments to be hardly more than mere guesses, there was usually little or no loss in accuracy. Our judgments are often based on unconscious cues; we react successfully, we know not how.’ None of the subjects with higher gains thought they were merely guessing. But then neither did a good number of the subjects with zero gain. Still, if Swenson is correct, then what he is describing is a form of ‘blindsight’: an ability to judge distance in the absence of any apparent sensory experience.

If this is correct, a further question is whether this ability to judge distance is an artefact of using pointing as a reporting mechanism? If we can converge onto a hidden hand, perhaps we can also achieve the opposite: pointing to the location of our convergence without any visual input. This concern, especially in light of the suggestion that ‘vision for action’ is broader than ‘vision for perception’ (Goodale & Milner, 1992), was the reason that manual pointing was traditionally regarded as unreliable for perceptual reporting (see Bingham et al., 2001).

5. Vergence-cognition linkage: For a number of my subjects, their response seemed more than guessing, but less than visual experience. As BF observed: ‘How you feel the distances, I think it’s very hard to describe.’ Which raises the question, could vergence be directly linked to our cognition of distance? Either via a pure judgement, or something more akin to an emotional response (a ‘sense’ of something close)? The vergence tonic cells and the vergence burst-tonic cells that report vergence angle in the midbrain (supraoculomotor area) feed into the cerebellum (dorsal vermis, caudal fastigial nucleus, and posterior interposed nucleus: see Leigh & Zee, 2015). But the extent to which the cerebellum is involved in cognition, rather than merely muscular control, continues to be debated (see Glickstein, 2007; Buckner, 2013 for conflicting accounts). In any case, vergence signals are also implicated in number of cognitive areas (such as the posterior parietal cortex: see Genovesio & Ferraina, 2004; and dorsolateral prefrontal cortex: see Alkan, Biswal, & Alvarez, 2011).

## 6 Vergence as a Relative Depth Cue

I am not the first to suggest that vergence is an ineffective cue to distance. Indeed, the idea that diplopia might explain vergence distance estimates is almost as old as the experimental study of vergence itself: see Hillebrand (1894)’s critique of Wundt (1862).

A couple of papers have since suggested that vergence is a relative, rather than an absolute, distance cue. Von Hofsten (1976) makes this point, rather paradoxically, by observing that subjects are very effective at estimating absolute distances but not changes in relative distance. Brenner & van Damme (1998) come to the same conclusion, but for exactly the opposite reason: they assume that subjects are poor at estimating absolute distance, but find evidence that subjects are actually quite good at relative depth judgements.

Why Brenner & van Damme (1998) assume absolute distance judgements are poor is unclear, even given the state of the literature in 1998 (see Appendix A). A number of the papers they cite investigate distances outside vergence’s effective range (>2m) or indirect measures (motion parallax, disparity scaling). And Brenner clearly regards vergence as an effective distance cue in more recent work: see e.g. Brenner & Smeets (2000) and de la Malla, Buiteman, Otters, Smeets, & Brenner (2016). But it is true that the literature at the time tended to show something close to a gain of 1, but with a consistent overshoot, typically of about 20cm. The choice appeared to be either endorse vergence as an absolute distance cue, but at the cost of its veridicality (Foley, 1980), or preserve its veridicality, but at the cost of limiting it to relative depth (Brenner & van Damme, 1998).

Should we embrace the idea that even if vergence isn’t an absolute distance cue, it is still an effective *visual* cue to relative depth? I don’t think so. Building upon earlier work (Wright, 1951; Foley & Richards, 1972; Enright, 1991; Enright, 1996), Brenner & van Damme (1998) provide impressive evidence that vergence is implicated in our relative depth judgements (see also Taroyan, Buckley, Porrill, & Frisby, 2000; Backus & Matza-Brown, 2003). But if vergence were to function as a *visual* cue to relative depth, then we would expect to see vivid motion-in-depth as our eyes track a dot moving backwards and forwards in space. First, we don’t: see the discussion of Regan et al. (1986), Harris (2006), and Howard (2008), in Appendix B. Second, what little motion we do see can be accounted for by retinal slip. Instead, as I discuss in Appendix B, vergence merely appears to function as a non-visual *kinesthetic* cue in the context of motion-in-depth, much like the sudden eye rotations I identified as one of my three confounding cues in Section 2. This interpretation is entirely consistent with Brenner & van Damme (1998), who merely demonstrate that vergence has *a* role to play in relative depth judgements, but who are unwilling to quantify the exact extent of this role.

Finally, if we were to admit that vergence was a *visual* cue to relative depth, would this erode the claim that vergence does not contribute to our absolute distance estimates? Only if we had accurate knowledge about our prior vergence distance. This explains the disagreement between Von Hofsten (1976) and Brenner & van Damme (1998): Von Hofsten suggests our absolute distance judgements are so good in the laboratory because we know our dark vergence state, whilst Brenner & van Damme suggest they are so bad because we do not. But both admit that this debate is largely confined to the laboratory. In real life we would be stuck in a near-infinite regress is we tried to judge distance based on our initial dark vergence state.

## 7 Do we see scale?

There are three directions we could take the conclusion that vergence and accommodation are ineffective cues to absolute distance.

First, we might continue the argument that vergence is not just an ineffective *visual* cue to absolute distance, but also an ineffective *visual* cue to relative depth, in the context of the motion-in-depth literature. This is question is addressed in Appendix B.

Second, we might consider how the conclusion that vergence is an ineffective cue to distance might be reconciled with the literature on vergence as a scaling cue for both size and disparity. This question is addressed in Appendix C.

Third, if vergence and accommodation are ineffective distance cues, we can begin to question whether there are any effective visual cues to absolute distance? This is my focus in this section. Specifically, what if all our cues to distance turn out to be:

1. Ineffective (vergence, accommodation),
2. Merely relative (diplopia, angular size of unfamiliar objects), or
3. Merely cognitive (angular size of familiar objects)?

In what sense could it be claimed that the visual system extracts absolute distance information? And in what sense could it be claimed that the visual system enables us to scale the visual scene; i.e. differentiate a small object up close from a large object far away?

I have already addressed the ineffectiveness of (a) vergence and (b) accommodation, so let me focus on (c) the angular size of unfamiliar objects, (d) the angular size of familiar objects, (e) motion parallax, and (f) vertical disparities.

Turning, first, to the angular size of *unfamiliar* objects, this is exactly what we want the visual system to disambiguate for us. And yet when Tresilian et al. (1999) asked subjects to estimate the distance of a luminous 0.7cm x 3cm piece of tubing based on angular size alone (monocular viewing through a 1.5 mm pinhole), they found a relationship of *y* = 0.73*x* + 11.77 for distances 30cm-100cm. Indeed, the addition of vergence and accomodation (binocular viewing, no pinhole) only marginally improved performance to *y* = 0.86*x* + 11.6. Similarly, when Mon-Williams & Tresilian (2000) tested accommodation as a distance cue for distances between 17cm-40cm, they found responses ranging from *y* = –0.30*x* + 4.79 to *y* = 0.62*x* + 1.55, but as soon as a size cue was introduced, performance ranged from *y* = 0.89*x –* 0.40 to *y* = 1.58*x –* 1.72. On this basis we should be confident that the angular size of unfamiliar objects is one of our most powerful absolute distance cues, even though it doesn’t provide us any absolute distance information. This finding has also been demonstrated by Lugtigheid & Welchman (2010); Sousa, Brenner, & Smeets (2011); and Sousa, Smeets, & Brenner (2012).

Recently de la Malla et al. (2016) came to the same conclusion when they tried to isolate motion parallax as an absolute distance cue. They found that motion parallax (a) had virtually no effect on the distance estimates of an isolated object, (b) had virtually no effect, even with the introduction of relative motion parallax cues, on binocular distance estimates, and (c) had only a negligible effect (gain of 0.115) on monocular distance estimates with the introduction of relative motion parallax cues. In the monocular condition, it was unclear what else could have been driving their subjects’ distance judgements apart from the angular size of the unfamiliar stimulus, leading de la Malla et al. (2016) to conclude that the angular size of unfamiliar objects must be one of our most important cues to absolute distance even though it doesn’t provide us with any absolute distance information.

By contrast, the angular size of *familiar* objects clearly does provide us with absolute distance information. But here the question is whether this familiarity functions as a *visual* cue to distance (as al-Haytham, c.1021 and Helmholtz, 1866 maintained), or merely as a *cognitive* cue that enables us to attribute distance to what we see (as Descartes, 1637 believed)?

Note that by *cognitive* I do not mean to imply that subjects have to consciously infer distance on the basis of familiar size. Instead, the inferences of ‘visual cognition’ can be *automatic* (we don’t have to do anything), *unconscious* (we are unaware of them), and *involuntary* (we cannot overrule them), and still be *post-perceptual*: see Linton (2017) and Linton (2018) in response to Cavanagh (2011) and Firestone & Scholl (2016). An example is reading: we do not have to *consciously* attribute meaning to words on a page, and yet it is hard to maintain that someone who understands a language and someone who does not have a different *visual* experience of the page itself.

I agree with the current consensus which appears to favour treating familiar size merely as a cognitive cue to distance. Specifically, it is not a source of information that the visual system uses to scale the scene: see Gogel (1969); Gogel (1976); Gogel & Da Silva (1987); Predebon (1992a); Predebon (1992b); Predebon, (1993); Predebon (1994); Predebon & Woolley (1994); Gogel (1998); Vishwanath (2014). For instance, Gogel (1976) found that familiar objects do not have the perceived motion parallax one would predict from subject head motion had familiar size determined their distance, whilst Predebon (1992b) found that the influence of familiar size could be vastly reduced simply by asking subjects for the apparent rather than physical size of the stimulus. It is also worth noting just how easily familiar size cues can be negated; for instance, in an Ames Room, or in the miniaturisation effect discussed in the context of Fig.11 and Fig.12 below.

**Fig.11.** Twin Stop 3D (2009) by Sasha Becher. © Sasha Becher. Used with permission from https://www.fotocommunity.com/photo/twin-stop-3d-anaglyph-sasha-becher/16025031 For more images see https://www.fotocommunity.com/user_photos/789955

**Fig.12.** Fig.12. Traffic on the Autobahn (2011) by Sasha Becher. © Sasha Becher. Used with permission from https://www.flickr.com/photos/stereotron/6603863175/in/album-72157673910516235/ For more images see https://www.flickr.com/photos/stereotron/

But the most appropriate demonstration of familiar size as a purely cognitive cue has not been conducted, namely turning an unfamiliar object (a luminous rectangle or a luminous circle) into a familiar object (a playing card or a coin) by means of a ‘Pepper’s ghost’ illusion that places them at the same optical distance. Replacing one (unfamiliar) object with another (familiar) one clearly gives us new distance information. The question is how this new distance information is processed? Does the new (familiar) object move forwards or backwards in space as it is relocated along the z-axis by the visual system? Or does it remain at the same position along the z-axis as the old (unfamiliar) object, only now the observer has a better understanding of what this distance corresponds to? If it is merely the latter, then familiar size is not functioning as a visual cue; it is not being used to locate objects in the 3D visual scene.

But if vergence, accommodation, and motion parallax have been shown to be ineffective, whilst familiar size has been shown to be merely cognitive, what else does this leave? Only really the ground-plane (Gibson, 1947; Gibson, 1950; Ooi, Wu, & He, 2001; Li & Durgin, 2012; Tyler, 2018), vertical disparities (Longuet-Higgins, 1981; Mayhew & Longuet-Higgins, 1982), and linear blur gradients (as evident in tilt-shift photography: Vishwanath, 2007; Held, Cooper, O’Brien, & Banks, 2009; Vishwanath & Blaser, 2010; Held, Cooper, O’Brien, & Banks, 2010). But all three of these cues are extremely limited in application. For instance, how often do you actually view an extended ground-plane in your daily life? And yet Gibson (1950) would have us believe ‘*that there is literally no such thing as a perception of space without the perception of a continuous background surface*.’ (And if that surface is not the ground-plane, what exactly gives it the scale by which to scale the rest of the scene?).

My inclination is to suggest that these, too, can be reframed as merely cognitive cues. Since this argument is most difficult to make in the context of vertical disparities, let me outline it before concluding. As far as I am aware there is only one paper that provides any evidence that vertical disparities scale the distance of a single object viewed in isolation: a single study in Appendix A of Rogers & Bradshaw (1995). But the reason this study is in an appendix, and not the main paper, is that the whole purpose of the main paper is to demonstrate that vertical disparities can distort the shape of a fronto-parallel plane, turning it into a 3D surface vividly curved in stereo depth. And this leads to two concerns:

First, vertical disparities appear to be double counted. Vertical disparities could either indicate (a) a flat object up close, or (b) a curved object further away, but in Rogers & Bradshaw (1995) they appear to be interpreted as both. Interestingly, the same concern arises in the context of tilt-shift miniaturisation, where the addition a linear blur gradient could either indicate (a) a reduction in distance, or (b) an increase in slant, but again is interpreted as both: see Vishwanath & Blaser (2010).

Second, the fact that the reduction in distance from vertical disparities in Rogers & Bradshaw (1995) is so closely associated with an increase in curved stereo depth suggests an alternative explanation, namely a cognitive association between (a) vivid stereo depth and (b) closer distances (reflecting our experience of an environment where disparity falls-off with distance^2^). This would suggest that we should experience the same reduction in apparent distance if horizontal disparities are increased without the introduction of vertical disparities. And this, indeed, is what we appear to find: Consider Fig.11 and Fig.12. The anaglyph format is useful because we can identify any significant vertical disparities in the scene simply by looking at the images without 3D glasses. There no significant vertical disparities in Fig.11 and Fig.12. Certainly nowhere near the magnitude that Rogers & Bradshaw (1995) suggest are required for vertical disparities to be effective. And yet, the miniaturisation effect is still strongly apparent in both images.

But if a cognitive association between accentuated stereo-depth and reduced distance explains the miniaturisation in Fig.11 and Fig.12, then it provides an alternative explanation for the results in the result in Appendix A of Rogers & Bradshaw (1995). The reduced distance from vertical disparities would be better understood as an inversion of the relationship suggested by Johnston (1991): it is accentuated or reduced 3D shape that scales distance, rather than the other way around. This is not to deny that vertical disparities make a contribution to our impression of distance, only that they make this contribution at the level of cognition (via accentuated stereo depth), and are no different from horizontal disparities in this regard.

The same miniaturisation effect can be achieved in real-world viewing using a telestereoscope, which uses mirrors to effectively increase the inter-pupillary distance. Helmholtz (1866) suggested that it ‘seems as if the observer were looking not at the natural landscape itself, but at a very exquisite and exact model of it, reduced in scale.’ Helmholtz (1866) attributes this effect to vergence scaling. Rogers (2011) to vergence scaling and vertical disparities. But we have already addressed Rogers’ vertical disparity explanation in the context of Fig.11 and Fig.12, whilst Helmholtz’s vergence explanation is subject to three key concerns:

First, it over-predicts the phenomenon: We do not get an impression of miniaturisation when we look at a photograph with near vergence.

Second, it under-predicts the phenomenon: The effects of telestereoscopic viewing can be experienced over hundreds of metres (see Sacks, 2010, on viewing St Paul’s Cathedral) which, even taking into account the change in the optical geometry by the increase in inter-pupillary distance, would still place the objects outside vergence’s effective range of 2m.

Third, Priot, Laboissière, Sillan, Roumes, & Prablanc (2010) find that when vergence is isolated in telestereoscopic viewing, subjects adapt their vergence rather than report a miniaturisation of visual space (for Priot’s further investigations of telestereoscopic adaption see Priot, Laboissière, Plantier, Prablanc, & Roumes, 2011; Priot et al., 2012; Priot, Neveu, Philippe, & Roumes, 2012; Priot et al., 2015; Priot, Vacher, Vienne, Neveu, & Roumes, 2018).

Indeed, given the miniaturisation effect in telestereoscopic viewing cannot be dismissed, what we appear to have in telestereoscopic viewing is a fundamental inconsistency between (a) vergence distance judgements (Priot et al., 2010) and (b) perceived scaling (Helmholtz, 1866; Rogers, 2011), suggesting that not only does vergence not explain the miniaturisation effect, it also doesn’t appear to function as an effective disparity scaling cue in the context of telestereoscopic viewing (on disparity scaling, see Appendix C).

The fact that this miniaturisation effect is ignored in some experimental conditions but not others leads Glennerster to a task-specific theory of visual space (see Glennerster, Tcheang, Gilson, Fitzgibbon, & Parker, 2006, to which Rogers, 2011 responds; see also Svarverud, Gilson, & Glennerster, 2012; Glennerster & Stazicker, 2017). Wagner comes to a similar conclusion in the context of familiar size (see Wagner, 2006; Wagner, 2012; Wagner & Gambino, 2016). For my part, I don’t believe that our perception of visual space is contingent upon the tasks we perform. Instead, many of the apparent inconsistencies in the literature reflect a distinction between *perception* and *cognition*: see Linton (2017) and Linton (2018). But this debate reflects a fundamental distinction that runs through the entire literature on visual space perception and is broader than the specific question of scale.

The final question is this: If all our visual cues to distance are either ineffective, relative, or merely cognitive, what kind of an account of 3D vision does this leave us with? One commenter worried it would leave us with an account where every object of fixation was seen at the same distance. Whilst Gogel (1969)’s ‘specific distance tendency’ doesn’t come to this extreme conclusion, he did argue that perception is at least biased in this direction. But I think this is a mistake. Indeed, it is a ‘category mistake’ (on ‘category mistakes’ see Ryle, 1949). If the visual system has no access to absolute distance information, the solution is not to posit a default distance (Gogel, 1969’s ‘specific distance tendency’ of 1.2–3.6m), and to suggest that the visual system uses *this* distance to scale the visual scene. Instead, the solution is to recognise that the visual system uses *no* distance to scale the visual scene. Indeed, there is no scaling of the visual scene by the visual system at all.

What is true under my account is that all objects of fixation are seen at the same *phenomenal depth*: the same extension along the z-axis in *phenomenal* space away from the observer. But this *phenomenal depth* has no *physical distance* intrinsically attributed to it: it can correspond to 20cm, or it can correspond to 20km. This coheres with the experience of many of the subjects in my experiments, who said that the dot was obviously some distance away, but they couldn’t tell how much: it could be close, or it could be miles away.

As I argue in Linton (2018), we experience something close to this effect whenever we watch a movie. The movie or tv screen is always at the same fixed phenomenal depth: the screen never appears to move towards us or away from us. And yet the fact that the movie is always at the same fixed phenomenal depth in no way impedes our enjoyment of the film: the movie can transition from a wide-panning shot of a cityscape to a close-up of a face without this transition being jarring. So attributing a variety of different scales to scenes viewed at a single fixed phenomenal depth appears to be something that comes naturally to humans.

Another context where I would argue that this principle applies is Emmert’s Law. Emmert’s Law suggests that the distance of an after-image varies with the distance of the object of fixation in the scene. The idea is that the after-image jumps from one distance to another as you change your fixation. But the problem is that Emmert’s Law applies monocularly, and at distances greater than 1m, so what could be scaling the distance of the after-image apart from cognitive cues such as familiar size? In which case, the ‘perceived’ change in the distance of the after-image could only be a post-perceptual cognitive effect.

Emmert’s Law is therefore open to an alternative explanation according to which it is the physical objects in the scene that jump forwards and backwards in *visual* depth, whilst the after-image remains at a fixed *visual* depth away from the observer. But because the real-world objects, unlike the after-image, have physical distances associated with them, as the physical objects jump forwards and backwards in *visual* depth, our *cognitive* attribution of distance to the fixed *visual* depth of the after-image will change, much like when we are watching a movie.

Finally, I am not the first to suggest that vision might operate without scale. Although Julesz (1995) ultimately came out in favour of an absolute depth account, he was at least willing to contemplate the possibility:

> ‘In the case of another qualia question (Do we sense absolute depth, or only relative depth?), the problem of the sensation of plasticity is as impenetrable as the essence of the sensation of ‘colour’…’
>
> ‘…I am still uncomfortable with the notion that absolute depth can be obtained by the mind…’
>
> ‘Except for echo-locating animals, can primates perceive absolute depth?’

It seems that we have finally come full-circle. Ptolemy’s ‘extramission’ theory of vision proposed scaling the angular size of objects using light rays that were emitted by the eyes and reflected back by objects. In practice some animals (bats, dolphins, whales, and even some birds and rodents) have evolved what is effectively an ‘extramission’ theory of audition to address this very concern. In the absence of echolocation in humans, Kepler (1604) and Descartes (1637) posited a number of visual cues that might plausibly have replaced Ptolemy’s ‘extramission’ theory of vision, most notably vergence and accommodation. But given their demonstrated ineffectiveness in my experiments, perhaps the time has come to admit the human visual system doesn’t extract absolute distance information from the environment, and to embrace a theory of vision without scale.

## Acknowledgements

This paper develops thoughts that were first articulated in Linton (2017) and Linton (2018).

This project was conducted at the Centre for Applied Vision Research, City, University of London as part of the author’s PhD research. This project, and the PhD as a whole, has been supervised by Christopher Tyler, and I would like to thank him for his advice on apparatus construction, stimulus presentation, and experimental protocol, as well as for our continued debates on the nature of visual scale (for his alternative account see Tyler, 2018). I would also like to thank Joshua Solomon, Simon Grant, Matteo Lisi, and Michael Morgan for their advice on apparatus construction, stimulus presentation, and experimental protocol, as well as for their continued discussion of the themes of this project. The experiments were conducted in Chris Hull’s ‘Ocular Optics Laboratory’ at City, University of London, and I would like to thank him for the provision of lab space and equipment for this project. I would also like to thank Matteo Lisi’s for his advice on statistical analysis, especially his help in applying lme4 and mclust5 in R. I would also like to thank John Barbur for testing the red filters for Experiment 2 in an autorefractor, and Ron Douglas for in testing them in a spectrophotometer.

I would like to thank Christopher Tyler and Simon Grant for their comments on the manuscript, Joshua Solomon for his comments on an earlier manuscript, and (in alphabetical order) Miriam Conway, Simon Grant, John Lawrenson, and Joshua Solomon for their comments when this work was presented at City, University of London (April 23 and June 8, 2018); Fulvio Domini, Julie Harris, Dhanraj Vishwanath, and Nick Wade for their comments when this work was presented at the Scottish Vision Group (Glencoe, March 16-18, 2018); and Wendy Adams, Kaan Akşit, Ben Backus, Marty Banks, Greg DeAngelis, Stephen Engel, Bas Rokers, Jannick Rolland, Laurie Wilcox, and Marina Zannoli for their comments when this work was presented at Frontiers in Virtual Reality (31^st^ CVS Symposium, University of Rochester, June 1-3, 2018).

Commercial Relationships: The author will be based at Facebook Reality Labs (formerly Oculus Research), Redmond, WA from September to December 2018.

## Appendix A Vergence as a Distance Cue

Whether vergence functions as an effective distance cue was one of the earliest questions of visual psychophysics. What is interesting is that many of those early debates continue to have relevance in the contemporary literature:

### a Vergence Micropsia and the Wallpaper Illusion

Meyer (1842) attributed the apparent reduction in size and distance in the wallpaper illusion to vergence. Today the wallpaper illusion continues to be cited in favour of vergence micropsia (see Lie, 1965; Hiroshi Ono, Mitson, & Seabrook, 1971), although Logvinenko et al. object on the basis that once the illusion is established, vergence can be increased by 2-3° without a change in apparent distance (Leont’ev, 1974; Logvinenko & Sokolskaya, 1975; Logvinenko, Nazarov, Sokolskaya, & Metcheryakov, 1980; Logvinenko & Belopolskii, 1994; Logvinenko, Epelboim, & Steinman, 2001; Logvinenko & Steinman, 2002). But as Kohly & Ono (2002) note, this objection (which has been well-recognised in the English-speaking literature since the mid-1980s: see Erkelens & Collewijn, 1985b; Regan, Erkelens, & Collewijn, 1986) rests on the assumption that vergence should be an effective cue to motion-in-depth despite a fixed angular size, and this question has been subject to its own long-running debate.

### b Dynamic Vergence and Motion-in-Depth

Wheatstone (1838)’s invention of the stereoscope enabled him to independently manipulate the vergence and the angular size of the stimulus. In Wheatstone (1852) he observed vergence micropsia, but no change in distance, as vergence was increased: ‘If we continue to observe the binocular picture whilst it apparently increases or decreases, in consequence of the inclination of the optic axes varying while the magnitude of the impression on the retinae remains the same, it does not appear either to approach or recede…’

This absence of motion-in-depth from vergence was confirmed by Erkelens & Collewijn (1985a; 1985b) when they tested 30° x 30° random-dot stereogram and found that (a) varying the vergence angle by 6° at 0.5Hz (maximum velocity 6°/s), and (b) varying the vergence angle by 5° at up to 1.5Hz (maximum velocity 6°/s to 13.5°/s depending on subject), had no effect on the perceived distance of the stereogram. The limits of this principle were illustrated by Regan, Erkelens, & Collewijn (1986), who found that although there is no motion-in-depth from vergence for (a) a 30° x 30° random-dot stereogram, nor for (b) a 33° x 38° field of dots, there is motion-in-depth from vergence for (c) a single 0.5° x 0.5° dot, and even (d) a single 1° x 0.2° bar. Motion-in-depth has also been reported from (e) a single 6.4° vertical line (Swanston & Gogel, 1986), (f) a single 2.1° x 2.1° spot (Howard, 2008), and even (g) a Siemens star viewed through an aperture (Howard, 2008; although the introduction of relative disparities might explain this).

Regan et al. (1986) describe these inconsistent results as a ‘paradox’. The explanation that has become accepted in the literature is that (a) vergence is a cue to motion-in-depth, but (b) it can be vetoed by the fixed angular size of larger stimuli: see Kohly & Ono (2002); Harris (2006); Nefs & Harris (2007; 2008); Howard (2008); Wismeijer & Erkelens (2009); Welchman, Harris, & Brenner (2009); González, Allison, Ono, & Vinnikov (2010); Allison (2013); Ono, González, & Lillakas (2014). This literature is re-evaluated in Appendix B.

### c Static Vergence and Near Distances

Reviewing the 19^th^ Century literature, Baird (1903) concluded that even ‘the most zealous advocates of the influence of convergence and accommodation have never maintained that this influence is operative save in the perception of near distances.’ Experiments over the last 60 years have only confirmed this hypothesis, finding very little effect beyond 2m (Collewijn & Erkelens, 1990; Howard, 2012). Given the geometry of vergence, this is unsurprising. As we see from Fig.13, there is relatively little change in the vergence angle beyond 1m.

**Fig.13.**
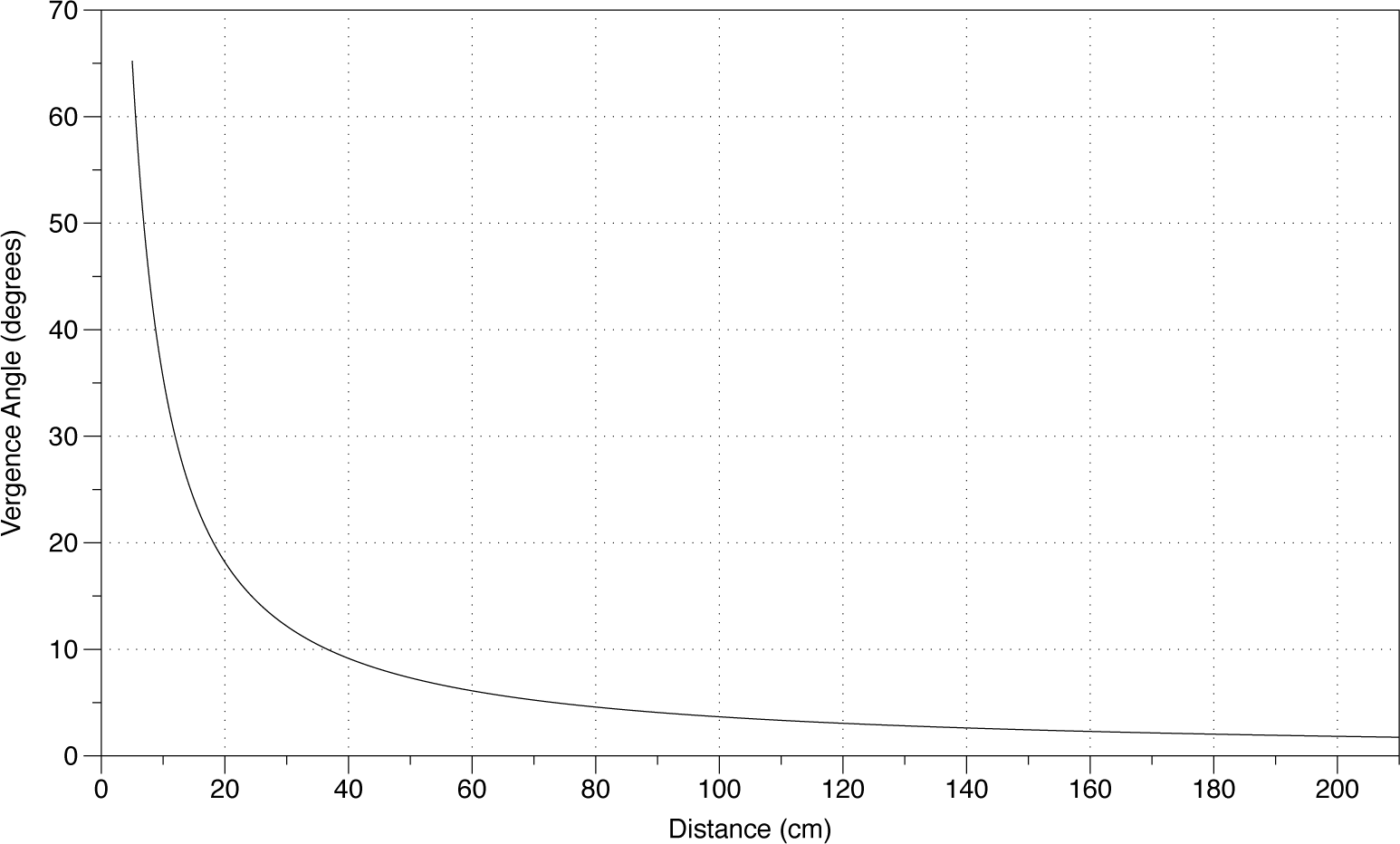
The fall-off of the combined vergence angles of the two eyes with fixation distance along the mid-line.

But studies within reaching space have provided us with some very strong results:

1. Swenson (1932) found that hidden-hand pointing to a small luminous disc was very accurate for distances between 25cm to 40cm, with an average error of less than 1cm.

2. Richards & Miller (1969) put the vergence of a point source in conflict with its size and luminance, but found that 2/3 of subjects were still able to set a reference to close to the vergence distance (32.75cm for a 28.5cm target) even though the target and the reference were not visible at the same time.

3. Foley & Held (1972) found that hidden-hand pointing to a point of light at distances between 15cm and 35cm was liable to produce a relationship of *y* = *x* + 25; i.e. on the whole subjects had a gain of 1, but they also overreached by a constant amount.

4. Gogel & Tietz (1973) was concerned with motion parallax, but as Foley (1977) reminds us they also asked subjects to estimate the distance of a point of light whilst their heads were stationary. Their results for 30cm (mean = 46cm, median = 28cm), and 91cm (mean = 100cm, median = 91cm), are actually much better than the ‘specific distance tendency’ for which the paper is remembered.

5. Komoda & Ono (1974) asked subjects to unravel a length of hidden rope in the x-axis to match the perceived distance of a 10° disc. At near distances (15cm to 60cm) they found distance gains of *y* = 0.34*x* + 19 and *y* = 0.37*x* + 19, but there was no full-cue condition to confirm the effectiveness of their rope-based reporting mechanism.

6. Von Hofsten (1976) asked subjects to indicate the distance of a point source on a hidden x-axis metre rule, and found a relationship of *y* = 0.92*x* + 21 for distances between 30cm to 61cm (or *y* = 0.96*x* + 20 if we just consider reaching space of 30cm to 46cm).

7. Foley (1977) found contradictory results for a point of light located 11cm to 39cm depending on which reporting mechanism was used. For verbal reports he found a relationship of *y* = 0.82*x* + 0.05, whilst for pointing *y* = 0.25*x* + 1.45. Full cue conditions were similarly inconsistent (*y* = 1.45*x –* 1.01 for verbal report, vs. *y* = 0.59*x* + 0.75 for pointing). Given pointing is pretty accurate in full-cue conditions (see below) this may reflect the fact that Foley had subjects pointing from below rather than from the side.

8. Foley (1980) suggested that all the distortions of binocular depth perception (specifically, 1. the lack of depth constancy in binocular stereopsis, 2. the curvature of the fronto-parallel plane, and 3. our inability to bisect distances) could be attributed to the misperception of distance from vergence. Whilst this paper helped to cement vergence as an absolute distance cue, it also implied that vergence was ‘non-veridical’, with a gain of only 0.5 (*y* = 0.5*x* + *c*). Although Foley didn’t advance any new data, this became the received wisdom for the next two decades.

9. Mon-Williams & Tresilian (1999) is the most influential study in recent years, and found a strong linear relationship between vergence and perceived distance for a point of light (*y* = 0.86*x* + 6.5; Fig.1) when subjects pointed with a hidden hand at distances between 20cm and 60cm. They also confirm the accuracy of this pointing method (*y* = 1.08*x –* 1.35) in full-cue conditions to defend it against Foley & Held (1972)’s and Foley (1977)’s suggestion that pointing is ineffective. Furthermore, Mon-Williams & Tresilian also contradict Foley (1980) by arguing that vergence is a veridical cue to distance (see Mon-Williams et al., 2001); albeit one that is subject to compression when studied in isolation due to cognitive decision strategies adopted in the context of uncertainty (subjects essentially hedge their bets: see Poulton, 1980).

10. Tresilian, Mon-Williams, & Kelly (1999) attempt to extend the range of their experiment (25cm to 100cm) by having subjects reach using a hidden pointer rather than a hidden hand. They find significantly reduced performance (*y* = 0.39*x* + 36.27). Interestingly, this can’t be atrtibuted to either (a) the increased range (since the near estimates are no more accurate than far ones), nor (b) the use of a pointer (since this gives a *y* = 0.99*x* + 0.2 relationship in full cue conditions). This result has largely been overlooked in the literature. It also demonstrates a worse performance than monocular distance estimates based on angular size alone (*y* = 0.73*x* + 11.77). Admittedly, the monocular angular size condition used a different stimulus (0.7cm x 3cm piece of tubing vs. the point of light used for vergence), but the addition of vergence and accommodation to the angular size stimulus only marginally improved performance (*y* = 0.86*x* + 11.6).

11. Mon-Williams, Tresilian, McIntosh, & Milner (2001) asked DF, the patient with visual form agnosia who formed the basis of Goodale & Milner (1992)’s ‘two-streams hypothesis’, and who cannot judge size cues, to participate in their pointing task in full cue conditions whilst her vergence was manipulated using prisms. Unlike ordinary subjects in full cue conditions, they found that DF’s pointing response reflected the manipulated vergence signal almost perfectly (*y* = 0.91*x* + 3.2) and accounted for 98% of the variance.

But it is also true that DF’s monocular reaches have distance scaling (see Carey, Dijkerman, & Milner, 1998), and given DF’s age (45) this seems unlikely to reflect either (a) accommodation or (b) vergence (via the vergence / accommodation linkage). Mon-Williams et al. (2001) attribute it to the vertical gaze angle (excluded in their experiment, so no challenge to their result). It might also reflect subtle motion parallax, vestibular cues, or even defocus blur given presbyopia, but again none of these challenge Mon-Williams et al. (2001)’s result.

12. Viguier, Clément, & Trotter (2001) presented subjects with a 0.57° disc at a distance between 20cm and 80cm for 5s, and then after 5s in darkness asked subjects to match the distance with a visible reference. They found subjects were close to veridical for 20cm, 30cm, and 40cm, but distances were increasingly underestimated beyond that: 60cm was judged to be 50cm, and 80cm was judged to be 56cm (see Fig.1).

The fall-off of performance in Viguier, Clément, & Trotter (2001) appears to challenge the hypothesis that vergence is a veridical cue to distance. In response, Scarfe & Hibbard (2017) have argued that subject performance in Viguier et al. (2001) might actually be optimal given noise in the vergence signal. This argument is an elaboration of Mon-Williams & Tresilian (1999) and Tresilian, Mon-Williams, & Kelly (1999)’s observation that symmetrical angular noise in the vergence signal will lead to an asymmetric range of possible distances. Mon-Williams & Tresilian (1999) and Tresilian, Mon-Williams, & Kelly (1999) argue that the fact the range is skewed towards further distances means that (a) the visual system will overestimate distance, and that (b) this overestimation is overcorrected for, leading to an underestimation. By contrast, Scarfe & Hibbard (2017) show that although the range is skewed towards further distances, this paradoxically leads to a reduction in the maximum likelihood.

Close to 40 years ago Foley (1980) observed: ‘Some writers held that the signal becomes inadequate for distant objects, but they attribute this to imprecision rather than inaccuracy.’ The contemporary literature therefore appears to have come full circle. (See also Brenner & Smeets, 2000’s observation that distance from vergence is as accurate as direction from version, once both have been articulated in angular terms). But the drop-off observed in Viguier et al. (2001) is not present in Mon-Williams & Tresilian (1999), suggesting that it is a consequence of Viguier et al. (2001)’s experimental paradigm, rather than inherent to vergence distance judgements. One possibility, suggested to me by Jannick Rolland, is that the constant angular size of the stimulus in Viguier et al. (2001) acts as a conflicting cue. Since Mon-Williams & Tresilian (1999) employ a point (with no discriminable angular size) there is no such cue-conflict, and this explains why I adopt their paradigm in my experiments.

13. Naceri, Chellali, & Hoinville (2011) has been influential in the virtual reality literature, where they test reaching for a ball of fixed angular size (8.89°) at distances between 25cm and 65cm. The results divided into two populations: 7 subjects with performance similar to Viguier et al. (2001), and 3 subjects with 0 gain. The performance of these 3 subjects did not improve in a full-cue virtual environment.

14. Swan, Singh, & Ellis (2015) tested subjects’ reaching responses to a virtual object viewed in an augmented reality set-up at distances between 34cm and 50cm. Because the stimuli were (a) 3D wireframe objects, and (b) reduced in size with distance, their results are not informative about vergence. However, their control test using a real object (*y* = 1.005*x –* 2.76) vindicates Mon-Williams & Tresilian (1999)’s pointing task.

15. Naceri, Moscatelli, & Chellali (2015) study distances outside our range (1.4m to 2.4m), but is notable for its methods: they use a robot to present 2 real-world luminous spheres with the same angular size (2.11°) at different distances for 1s each with a 1s interval between them. At these distances the JND was only 10cm ± 1cm, although these results are open to an alternative ‘sequential stereopsis’ interpretation (see Enright, 1996).

16. Campagnoli, Croom, & Domini (2017) question vergence’s veridicality. Like Foley (1980) and Johnston (1991) before them, they suggest that distortions of binocular stereopsis reflect distortions of distance. They tested a stimulus made up of 3 virtual bars (0.5cm wide and 6cm high) arranged in an isosceles triangle (with one rod 3cm-6cm in front of two rods 4cm apart) at two different distances (45cm and 55cm) and had subjects reach for them. Their results appear to be an improvement on Viguier et al. (2001) with 44cm for the 45cm stimulus, and 52cm for the 55cm stimulus, when the rods were separated 3cm in depth (this reduced to 43cm and 50cm respectively when the rods were separated 6cm in depth). But even though the study was intended as a test of vergence, the size of the stimulus varied with distance. When they tested a single rod, they found a linear relationship of *y* = 0.77*x* + 10.3 for distances between 36cm and 48cm, but again this might have been influenced by the changing angular size, as well as the inclusion of a backplane (see Magne & Coello, 2002).

Summarising these results, there seems to be no question in the contemporary literature that vergence is an effective egocentric distance cue. The only real question is whether it is veridical within reaching space or not?

## Appendix B Vergence as a Motion-in-Depth Cue

The conclusion that vergence is an ineffective visual depth cue would explain the absence of motion-in-depth in Erkelens & Collewijn (1985a; 1985b), and would help us to unify the distance and motion-in-depth literature. However, the finding in Erkelens & Collewijn (1985a; 1985b) has been heavily revised over the last 30 years, and there is now significant evidence that vergence contributes to motion-in-depth. Can my thesis explain these observations?

I would argue that it can. The raft of papers that emerged in 2006-2010 only sought to prove that vergence made *a* contribution (as opposed to *no* contribution) to our motion-in-depth estimates. In the words of Welchman et al. (2009), it was ‘surprising that the view that vergence velocity signals do not support perceptual estimation has remained largely unchallenged for over twenty years.’ By contrast, I am happy to admit that vergence makes a modest contribution to our motion-in-depth estimates. But my question is whether this contribution is *visual* (vergence affects our visual experience), or merely *kinesthetic* (a physical sensation that confirms / denies our visual experience)?

To give an illustrative example, consider how *visual* (optic flow) and *kinesthetic* (vestibular) cues interact in the experience of illusory self-motion. Vestibular cues are not necessary for illusory self-motion (Fischer & Kornmüller, 1930), e.g. when we see a neighbouring train pull away in a railway station whilst we are stationary. Nonetheless, the addition of vestibular acceleration can contribute to the illusion of self-motion (Ash, Palmisano, & Kim, 2011). But, and this is the crucial point, there is no suggestion that this contribution has to be *visual*: It is not as if the optic flow has to be seen as flowing faster in order to incorporate this vestibular cue into our determination of self-motion. Instead, the visual (optic flow) and the non-visual (vestibular) cues merely feed into a common post-perceptual determination. Indeed, as Ash, Palmisano, & Kim (2011) observe, there is no ‘mandatory fusion’ between the *visual* (optic flow) and *kinesthetic* (vestibular) cues, which is often taken to be the litmus test of a truly perceptual effect: see Hillis, Ernst, Banks, & Landy (2002).

But if this holds true for self-motion, why can’t it hold true for object motion? After all, the two scenarios are symmetrical: the same *visual* cue (optic flow, which is identical whether it is caused by self-motion or object motion) is confirmed by a *kinesthetic* cue (*vestibular* in the case of self-motion, *vergence* in the case of object motion) by feeding into a common post-perceptual determination. Let us consider how this hypothesis fits with the empirical literature:

1. The first argument against vergence as a purely *kinesthetic* cue is that the same change in angular rotation would produce the same apparent motion, even though the motion-in-depth for 0° to 5° (from the horizon to 73cm) is much greater than the motion-in-depth for 5° to 10° (from the 73cm to 37cm). But Lugtigheid, Brenner, & Welchman (2011) find that motion-in-depth from vergence isn’t scaled for absolute distance, and does in fact simply rely upon the change in the angular rotation of the eyes.

2. The second argument against a purely *kinesthetic* account of vergence is that such a cue would still be present in those instances where we don’t see motion-in-depth (for instance a 30° x 30° stereogram oscillating in depth: Erkelens & Collewijn, 1985a; 1985b), so why do we fail to detect motion in this context? The problem is starkly illustrated by Welchman, Harris, & Brenner (2009), who confirm that subjects can use vergence to judge the direction of motion of a 0.1° x 0.1° square, but fail for a 22° x 17° field of triangles: ‘This confirms previous findings that extra-retinal signals alone do not inform perceptual estimates of motion direction for large-field stimuli that do not change in retinal size (Erkelens & Collewijn, 1985b; Regan et al., 1986).’ As Welchman et al. explain, they used a field of triangles to ‘make sure that motion-in-depth of the large background was imperceptible.’ Both Welchman et al. and I agree that vergence is not functioning as a *visual* cue in this context.

But unlike Welchman et al., I argue that vergence must be acting as a non-visual *kinesthetic* cue in this context, since (contrary to the above quotation) subjects performed quite well when detecting the motion-in-depth of the 22° x 17° field of triangles. When we look at Welchman et al.’s data (their fig.3a, 0ms), we find thresholds for detecting the direction of motion for a 22° x 17° field of triangles are around 0.7°/s (12cm/s), which is (a) only double the threshold for a 0.1° x 0.1° square, and (b) an order of magnitude lower than the 6°/s to 13.5°/s tested by Erkelens & Collewijn (1985b) for which subjects are still *visually blind* to motion. So subjects are unable to see the motion, but they are nonetheless able to reliably judge its direction. This is only a paradox if we are committed to thinking that all motion-in-depth cues must operate *visually*. They clearly don’t in the context of illusory self-motion (with vestibular cues), so why think they must in the context of object motion?

But why has this apparent paradox been overlooked? Because Welchman et al. (2009)’s apparatus had a theoretical maximum of 2.2°/s (50cm/s), thresholds had to be less than 0.4°/s (7.5cm/s) to reach statistical significance. Since the threshold for the 22° x 17° field of triangles was ≈ 0.7°/s (12cm/s) it didn’t reach statistical significance, but the threshold for the 0.1° x 0.1° square (after 300ms) was ≈ 0.3°/s (6cm/s) so it did (cf. the performance after 100ms and 200ms). In the context of the 22° x 17° field of triangles Welchman et al. interpret the absence of statistical significance as confirmation of the null hypothesis. But this gives a misleading impression of the modest differences between their conditions (see their fig.3a). Specifically, it eradicates the performance in the 22° x 17° field of triangles condition, rather than explaining it. The only explanation for this performance, if we are committed to the idea that subjects are *blind* to the motion-in-depth of large-field stimuli, is that subjects were aware of their eye movements by means of a non-visual *kinesthetic* cue: even if subjects couldn’t see the motion, they could feel the rotation of their eyes.

3. The third argument against a purely *kinesthetic* account of vergence is that although *visual* motion-in-depth was absent in Erkelens & Collewijn (1985a; 1985b), it was later found to be present in Regan et al. (1986) when subjects viewed a single dot moving in depth.

But I would argue that this is actually the strongest argument in favour of vergence as a purely *kinesthetic* cue: If vergence were an effective *visual* cue to motion-in-depth, then we would expect a vivid impression of motion as the eyes tracked the single dot. And yet this isn’t the case: Regan et al. (1986) describe the motion-in-depth as ‘weak’, Harris (2006) ‘confirms that Z-direction motion is indeed difficult to detect’, and the subjects in Howard (2008) report only 3.7cm gain for a 37.5cm change in vergence. Indeed, the motion-in-depth was so poor that Harris (2006) concluded that ‘the visual system appears not to use vergence eye movement signals in the perception of the Z-component of 3-D motion.’ This is despite the fact that we know from Brenner & van Damme (1998) that vergence contributes to effective *judgements* of relative depth, so whatever contribution vergence is making, it doesn’t appear to be *visual*.

Furthermore, the modest motion-in-depth that is reported can be explained by retinal slip. Welchman et al. (2009) exclude the possibility that retinal slip is *solely* responsible for our motion-in-depth estimates, but that isn’t what’s being claimed. We’ve just seen that in Welchman et al. (2009) motion detection thresholds for a dot are only half those of an extended field. Consequently, we might expect half our motion detection to come from the non-visual vergence signal (that also accounts for the motion detection for the extended field), and half to come from the visually apparent retinal slip. This is consistent with Welchman et al. (2009)’s finding that both cues contribute relatively equally to motion detection.

Welchman et al. (2009) also find that subjects perform substantially worse in detecting the motion of a dot when they track its motion with vergence rather than simply keeping their eyes fixed. This suggests that whatever contribution vergence does make to the discrimination of motion-in-depth, it is substantially less than the retinal slip it replaces. In this regard, vergence also appears to supress motion-in-depth rather than being an effective cue to it.

4. The fourth argument in favour of vergence as a *visual* cue is that when motion-in-depth is produced by ‘looming’ (a sudden increase in angular size), the addition of a consistent vergence signal appears to accentuate the motion (Heuer, 1987; Brenner, Van Den Berg, & Van Damme, 1996; ‘texture alone’ in Howard, Fujii, & Allison, 2014, fig.4; cf. ‘texture alone’ in Howard et al., 2014, fig.5). However:

First, the size of this effect is relatively modest, if it is present at all: Brenner, Van Den Berg, & Van Damme (1996) find only ‘a slightly lower velocity’ from the absence of vergence, and in the one instance Howard, Fujii, & Allison (2014) do find an improvement, the gain is only about 10% (from 20cm of motion-in-depth to 22cm). There is no reason why an effect of this magnitude couldn’t be accounted for by a purely *kinesthetic* cue.

Second, there is virtually no evidence that vergence in the wrong direction has the opposite effect (Heuer, 1987; ‘texture alone’ in Howard et al., 2014, fig.5; cf. Swanston & Gogel, 1986), even though, as we have already noted, such reduced performance (‘mandatory fusion’) is the litmus-test for a truly perceptual effect: see Hillis et al. (2002). Indeed, Heuer (1987) suggests that subjects can consciously choose which of the two contradictory signals – vergence or looming – to attend to. So just as with illusory self-motion (Ash, Palmisano, & Kim, 2011), we find no ‘mandatory fusion’ between the *visual* and *kinesthetic* cues. Indeed, this appears to be a trend when *visual* cues are integrated with *physical sensations*, be they *kinesthetic* or (in the case of Hillis et al., 2002) *haptic*.

5. Fifth, Lugtigheid et al. (2011) find vergence can modulate motion-in-depth from relative disparity by up to 60%. But this impressive figure needs to be put into context: (1) The relative disparity is moving towards us at roughly 3.7cm/s. (2) Adding a convergent eye movement of 13.6cm/s adds an impression of forward motion of about 1cm/s, so the relative disparity in the stimulus only has to travel at 2.7cm/s to achieve the same motion. (3) Adding a divergent eye movement of 13.6cm/s seems to wipe off forward motion by about 1.3cm/s, so the relative disparity has to travel at 5cm/s to achieve the same forward motion. So, as with vergence’s contribution to looming, the gain is an order of magnitude less than the vergence signal itself. And again, there is no reason why this effect could not be accounted for by a *kinesthetic* cue.

6. Sixth, Nefs & Harris (2008) find vergence is important for the induced motion illusion, where a fixed target appears to move in the opposite direction from an actually moving target. But an alternative explanation for this finding is that induced motion in depth only occurs when (a) you are not fixated on the static target, and (b) there is relative disparity between the static and moving target. The only experimental condition in Nefs & Harris (2008) that satisfied those two conditions necessarily involves changing vergence.

In conclusion, over the last decade the motion-in-depth literature has been driven by the assumption that ‘the distance of a stationary object can be judged on the basis of vergence alone. So why was motion-in-depth not produced by changing vergence?’ (Howard, 2012). Once we realise that stationary vergence isn’t an effective distance cue, the impetus for this literature is removed. The last decade of effort has seen the accumulation of studies with very modest gains in motion-in-depth from vergence. This is consistent with (and indeed, would appear to confirm) the notion that changing vergence merely functions as a *kinesthetic* cue.

## Appendix C Vergence as a Scaling Cue

Vergence is supposed to be fundamental to two scaling mechanisms: (1) size constancy, and (2) depth (or 3D shape) constancy. These two scaling mechanisms have to be reassessed in light of vergence’s ineffectiveness as a distance cue:

1. Size Constancy: When we scale a visual scene, we are trying to differentiate a small object up close from a large object far away. In this sense, size and distance are inextricably linked. We’ve known since Ptolemy (c.160 AD) that the visual angle of the stimulus does not determine this question, and that the visual system has to rely on distance information to specify the size as well and distance of the stimulus. Some of the earliest studies of vergence as a distance cue (Meyer, 1842; Wheatstone, 1852) suggested that vergence scales the size of the stimulus, because the stimulus appears to shrink as vergence is increased. But if vergence is an ineffective distance cue, then how are we to explain this ‘vergence micropsia’?

One suggestion is that vergence is a distance cue for size and disparity, but not for distance itself. Ono & Comerford (1977) consider the possibility that ‘the question whether convergence serves as a cue to distance can be divided into two parts: (a) does oculomotor adjustment provide distance information for the visual system and (b) is the distance information provided by oculomotor adjustment, if any, used to make distance judgments?’ Bishop (1989) makes a similar point. But this would be a surprising conclusion to have to come to. First, it seems paradoxical that distance should be a cue for size and disparity, but not distance itself. Second, as we have just observed, size and distance are inextricably linked so far as ‘scale’ is concerned: we want to differentiate a small object up close from a large object far away, yet what we’d have is one parameter (size) changing without the other (distance).

The alternative is to explore whether vergence micropsia is an artefact of stimulus presentation. We can begin to see why it might be if we cross-fuse the two coins in Fig.14.

**Fig.14.**
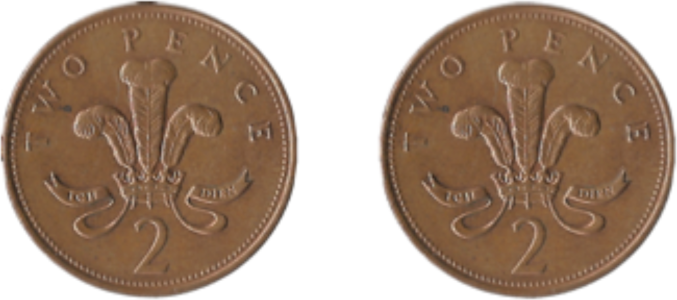
Two coins to cross-fuse as demonstration of vergence micropsia.

What we notice are two idiosyncrasies of vergence micropsia: First, vergence micropsia shrinks the central fused coin, but appears to leave the two monocular flankers unaffected (this works in the opposite direction when the coins are parallel-fused). Second, this asymmetry in size between the central fused coin and the monocular flankers is only accentuated as we reduce our distance to the screen.

What might explain this effect? Consider how the two coins are viewed when your eyes are not crossed: The distance of the left coin from the right eye is slightly greater than the distance of the right coin from the right eye. Each coin is made up of a larger retinal image (right eye in the case of the right coin, left eye in the case of the left coin) and a smaller retinal image (left eye in the case of the right coin, right eye in the case of the left coin). But when we cross-fuse, what happens? The central coin is made up to the two smaller retinal images, whilst the two monocular flankers are made up of the remaining larger retinal images, leading to the asymmetry that we see between the central fused coin and its monocular flankers.

The same concern applies when vergence is varied in a stereoscope by increasing the separation between the stimuli for the left and right eye: First, as the stimulus for each eye travels increasingly in the opposite direction, the distance of the stimulus from the eye increases, introducing small reductions in x-axis and y-axis size. Second, as the stimuli are presented on a fronto-parallel display, when the distance from the eye increases the stimuli are also rotated relative to the eye, introducing another small reduction in x-axis size. Third, this rotation doesn’t just reduce the x-axis size, it also distorts the shape of the object. We simply don’t know the x-and y-axis size of the eventual fused percept is affected when the visual system has to reconcile two inconsistently distorted shapes.

One option is to build these three distortions into the stimulus, and reduce them as vergence increases, thereby keeping the distortions constant with vergence, and see if vergence micropsia nonetheless persists. But it has also been hypothesised that a reduction in pupil size, brought on by the ‘near triad’ response, could impact retinal image size (Enright, 1975; although it is not clear why), so it would be better if we could control for this factor as well.

The alternative is to fix the retinal image with after-images viewed in darkness. There are reports of vergence micropsia when subjects view an after-image of their hand and then move their real hand backwards and forwards in space (or close variations of this paradigm; see: Taylor, 1941; Gregory, Wallace, & Campbell, 1959; Morrison & Whiteside, 1984; Suzuki, 1986; Carey & Allan, 1996; Mon-Williams et al., 1997; Bross, 2000; Ramsay, Carey, & Jackson, 2007; Sperandio, Kaderali, Chouinard, Frey, & Goodale, 2013; Zenkin & Petrov, 2015; see also Niechwiej-Szwedo & Steinbach, 2007 vs. Niechwiej-Szwedo et al., 2007). But there are two concerns with this literature:

First, all these papers describe vergence micropsia as an application of Emmert’s Law, according to which the physical size of an after-image varies with its physical distance (Emmert, 1881; Darwin, 1786). But if this is correct, then vergence is not acting as a size cue over and above its acting as a distance cue: vergence is neither (a) changing the apparent angular size of the after-image, nor (b) is there a reduction in apparent physical size divorced from a reduction in apparent physical distance (see Zenkin & Petrov, 2015; Chen et al., 2018).

This is at odds with the traditional vergence micropsia literature, according to which increasing vergence causes a reduction in apparent angular size (McCready, 1965). Indeed, vergence micropsia is often yoked together with the moon illusion as an instance of a ‘visual angle illusion’ (Enright, 1975; McCready, 1985; Suzuki, 2007). But if ‘vergence micropsia’ just means ‘an application of Emmert’s Law to vergence as a distance cue’ then, contrary to this literature, there is no vergence-specific size-scaling phenomenon to analyse over and above vergence as a distance cue. Lou (2007) claims to have found that increased vergence reduces the angular size of after-images, but the task (matching the size of a reference to the size of an after-image in full-cue conditions) introduces the very physical size cues that using after-images were meant to control for. Indeed, if the results are interpreted in terms of physical rather than angular size, Lou (2007) is evidence of vergence *macropsia* not *micropsia*.

Second, does this literature establish vergence as the distance cue in question? I don’t believe so: First, if vergence were the distance cue in question, then the after-image literature would be inconsistent with the motion-in-depth literature. Both vary vergence as subjects view a stimulus of fixed angular size. In the after-image literature, vergence wins out and the change in distance modulates the apparent size. But in the motion-in-depth literature, the fixed angular size wins out and vetoes an apparent change in distance. Second, in many of the experiments hand movements could be providing the requisite change in distance information. Mon-Williams et al. (1997) try to exclude this by having subjects track a moving LED between 20cm and 50cm, but this brings its own concerns (change in size of the LED, diplopia from retinal slip, or, as Ramsay et al., 2007 note, the height of the LED).

This literature begs a more general question about the integration of vision and proprioception. The rubber-hand illusion (Botvinick & Cohen, 1998) provides a vivid illustration of just how easily these cues can come apart. If (a) my hypothesis is correct, and vision doesn’t provide us with absolute distance information, but (b) touch does, then perhaps the fact that vision and touch provide us with very different metric depth information helps to explain why a further level of integration is required, and how these cues can come apart. For further discussions of these issues see Linton (manuscript).

2. Shape Constancy: Mis-scaled binocular disparities are liable to distort both the 3D shape of objects and the 3D shape of the visual field itself. To avoid this, the visual system is thought to employ a second scaling mechanism. Although traditionally referred to as ‘depth constancy’, I prefer ‘shape constancy’ because it brings out what is genuinely at stake:

a. Shape of Objects: Whilst angular size varies in proportion to distance, binocular disparity varies in proportion to distance^2^. What this means is that the z-axis depth of an object drops off more rapidly with distance than its x-axis and y-axis size, so horizontal disparity does not preserve the 3D shape of an object as its distance increases. Whilst this coheres with our impression of real-world scenes extending off into the far distance (the z-axis depth of distant buildings appears to fall off more quickly than their size, leaving them, in Vishwanath, 2010’s terms, ‘almost *pictorial’*), for a couple of decades it was believed that within vergence’s effective range (up to 2m) the visual system not only compensated for this drop-off in z-axis depth relative to x-axis and y-axis size (i.e. preserved the object’s 3D shape), but actually recovered the object’s absolute metric depth: see Wallach & Zuckerman (1963); Fried (1973); Ono & Comerford (1977); Wallach, Gillam, & Cardillo (1979).

This consensus was eroded by Foley (1980) and Johnston (1991), the latter finding that the 3D shape of a cylinder defined by disparity was distorted with distance (at 53.5cm it was elongated towards the viewer, at 107cm it was veridical, and at 214cm it was compressed). This distortion of 3D shape with viewing distance has been confirmed by Glennerster, Rogers, & Bradshaw (1996); Bradshaw, Parton, & Glennerster (2000); and Scarfe & Hibbard (2013). There have been four responses to this shape inconstancy: (1) Quarantine it to specific tasks / contexts (Glennerster et al., 1996; Bradshaw et al., 2000). (2) Eradicate it by introducing other depth cues (Scarfe & Hibbard, 2013; Guan & Banks, 2016). (3) Suggest that stereo-depth and 3D shape are two different things (Hibbard, 2008; Vishwanath, 2010; Vishwanath, 2014). (4) Simply admit that the 3D shape of objects can be distorted with viewing distance (Morgan, 1989; Johnston, 1991; Campagnoli et al., 2017).

But the admission that 3D shape can be distorted with viewing distance only addresses the fact that shape constancy is not veridical. I still need to explain why vergence appears to make some contribution to shape constancy. The typical suggestion is that vergence is scaling disparity, but using a non-veridical distance estimate (Foley, 1980; Johnston, 1991; Campagnoli et al., 2017). But there is another phenomenon that needs explaining: When we increase our distance from a stereo-image, by walking away or moving our head back, the scene appears to expand in z-axis depth relative to the x-axis and y-axis, and compress when we approach it: try it with Fig.11 and Fig.12. There are two things to notice: First, the geometry of the scene is vividly distorted as we move our head back and forth, suggesting (vs. Scarfe & Hibbard, 2013) that the addition of other depth cues does not eradicate such distortions. Second, the scaling of the depth in the scene is the opposite of the relationship we would expect: as vergence reduces, the depth in the scene increases.

This observation is explained away by Wallach et al. (1979) who argue that for a fixed-disparity image (such as a stereo-image) disparity falls off linearly with distance, so depth scaling (which responds to a fall-off of disparity with distance^2^) will necessarily overcompensate. However, there is a problem. When disparity is fixed, and vergence is increased, this is supposed to increase, not decrease, the depth from disparity. Indeed, this is the classic claim of the literature: see Helmholtz (1866) on telestereoscopic viewing, or more recently Cumming, Johnston, & Parker (1991).

But there is another explanation. What is common to both scenarios is a reduction in x-axis and y-axis size, due to an increase in physical distance (in Wallach et al., 1979) or vergence micropsia (in Cumming et al., 1991). Perhaps the increased impression of depth in both of these contexts is simply a function of constant retinal disparity seen in relation to a reduction in x-axis and y-axis size? Properly testing this hypothesis would require a manipulation of vergence in my apparatus whilst subjects viewed stereoscopic after-images of the kind explored by Wheatstone (1838); Dove (1841); and Lugtigheid et al. (2014).

b. Shape of the Visual Field: The binocular disparity of a fronto-parallel surface varies with eye rotation, and therefore vergence distance. But the shape of fronto-parallels do not appear to change with distance, which suggests a scaling mechanism. Fronto-parallel constancy was originally seen as less effective than shape constancy (see Ono & Comerford, 1977’s discussion of Luneburg, 1947). Then they were regarded as equally non-veridical (Foley, 1980). Now this position has reversed, and it is fronto-parallel constancy which is regarded as close to veridical, and shape constancy which is regarded as comparatively ineffective (Rogers, 2006). Why there should be these discrepancies in the literature, if vergence is an effective cue to distance, is far from a settled question. Furthermore, the fact that fronto-parallel surfaces don’t show shape constancy from vertical disparities (Rogers & Bradshaw, 1995), even though vertical disparities are largely a product of eye rotation, is worthy of attention.

